# Experimenter sex modulates mouse biobehavioural and pharmacological responses

**DOI:** 10.1101/2022.01.09.475572

**Authors:** Polymnia Georgiou, Panos Zanos, Ta-Chung M. Mou, Xiaoxian An, Danielle M. Gerhard, Dilyan I. Dryanovski, Liam E. Potter, Jaclyn N. Highland, Carleigh E. Jenne, Brent W. Stewart, Katherine Pultorak, Peixiong Yuan, Chris F. Powels, Jacqueline Lovett, Edna F. Pereira, Sarah M. Clark, Leonardo H. Tonelli, Ruin Moaddel, Carlos A. Zarate, Ronald S. Duman, Scott M. Thompson, Todd D. Gould

## Abstract

Differential rodent responses to the sex of human experimenters could have far reaching consequences in preclinical studies. Here, we show that the sex of human experimenters affects mouse behaviours and responses to the rapid-acting antidepressant ketamine and its bioactive metabolite (*2R,6R*)-hydroxynorketamine. We found that mice manifest aversion to human male odours, preference to female odours, and increased susceptibility to stress when handled by male experimenters. This male induced aversion and stress susceptibility is mediated by the activation of brain corticotropin-releasing factor (CRF) neurons projecting from the entorhinal cortrex to hippocampal area CA1. We further establish that exposure to male scent prior to ketamine administration activates CRF neurons projecting from the entorhinal cortex to hippocampus, and that CRF is necessary and sufficient for ketamine’s *in vivo* and *in vitro* actions. Further understanding of the specific and quantitative contributions of the sex of human experimenters to different experimental outcomes in rodents may lead not only to reduced heterogeneity between studies, but also increased capability to uncover novel biological mechanisms.

## Main text

Lack of replicability of experimental results in different laboratories may be due to unrecognised experimental variables that are not appropriately controlled ^1^. The sex of the human experimenter is rarely considered as a biological variable that can affect experimental results and is usually neither accounted for by design or statistical methodology, nor reported in experimental methods ^2–4^. However, there is evidence rodents may be able to differentiate the sex of human experimenters, and this discrimination could have measurable effects on their behavioural and/or biological responses. Indeed, exposure of rodents to the scent of male, but not female, experimenters was shown to increase anxiety-related behaviours and stress-induced analgesia . Here, we investigated the role of the experimenter’s sex on stress-induced maladaptive behaviours and how sexually dimorphic scents might affect rodent biobehavioural responses to pharmacological antidepressant treatments (see *Supplementary Table 1* for details on experimental designs, statistical tests used and *n* numbers).

### Human experimenter scent modulates behavioural responses

We first assessed whether CD1, BALB/c, and C57BL/6J mice prefer to interact with a cotton swab rubbed across the skin (*cubital fossa*, inner wrist and mastoid) of human males compared with human females. Mice of both sexes consistently preferred the female-scented swabs (Fig. 1A), while zinc sulphate-induced anosmia abolished this preference (Fig. 1B), suggesting that these effects are smell-driven. Also, when given the choice between a swab dampened with water *vs* human male or female skin-exposed swabs, mice showed aversion to the male swabs, whereas they displayed preference for the female swabs (Fig. 1C). In addition, mice manifested avoidance to male-worn t-shirts, whereas they exibited preference to female-worn t-shirts in the real-time place preference paradigm (Fig. 1D). Moreover, to directly compare the behavioral response of the mice to male, female and control scent, we used a three-arm Y-maze. We attached one swab within each arm and measured the time spent in each arm. In agreement with Fig 1A-D outcomes, we found that mice spent less time in the male swab containing arm compared with either the control or female experimenter scents (Fig 1E,F). No difference was observed between control and female expereiimenters scents (Fig 1E,F). To test the specific aversive/rewarding properties of male *vs* female experimenter scent, mice were assessed for conditioned-place preference/aversion, where mice were exposed to distinct scents within each of two distinct compartments and subsequently assessed if any conditioned preference developed. Male experimenters’ scent induced place aversion (Fig. 1 G,H), supporting an aversive aspect to male scent; however, neither place preference nor aversion was observed following exposure to female experimenters’ scent (Fig. 1 G,H). We performed additional analyses of these prior experiments (Fig. 1A-H) using receiver operating characteristic (ROC) curves. A ROC curve is a commonly used analysis for binary classifiers, in our case male or female/control experimenters, to demonstrate if the effects are most like true or false. We found that ROC curve is robustly above the chance line, indicating that effect of male experimenters is a true positive (Fig 1I).

**Fig. 1:**
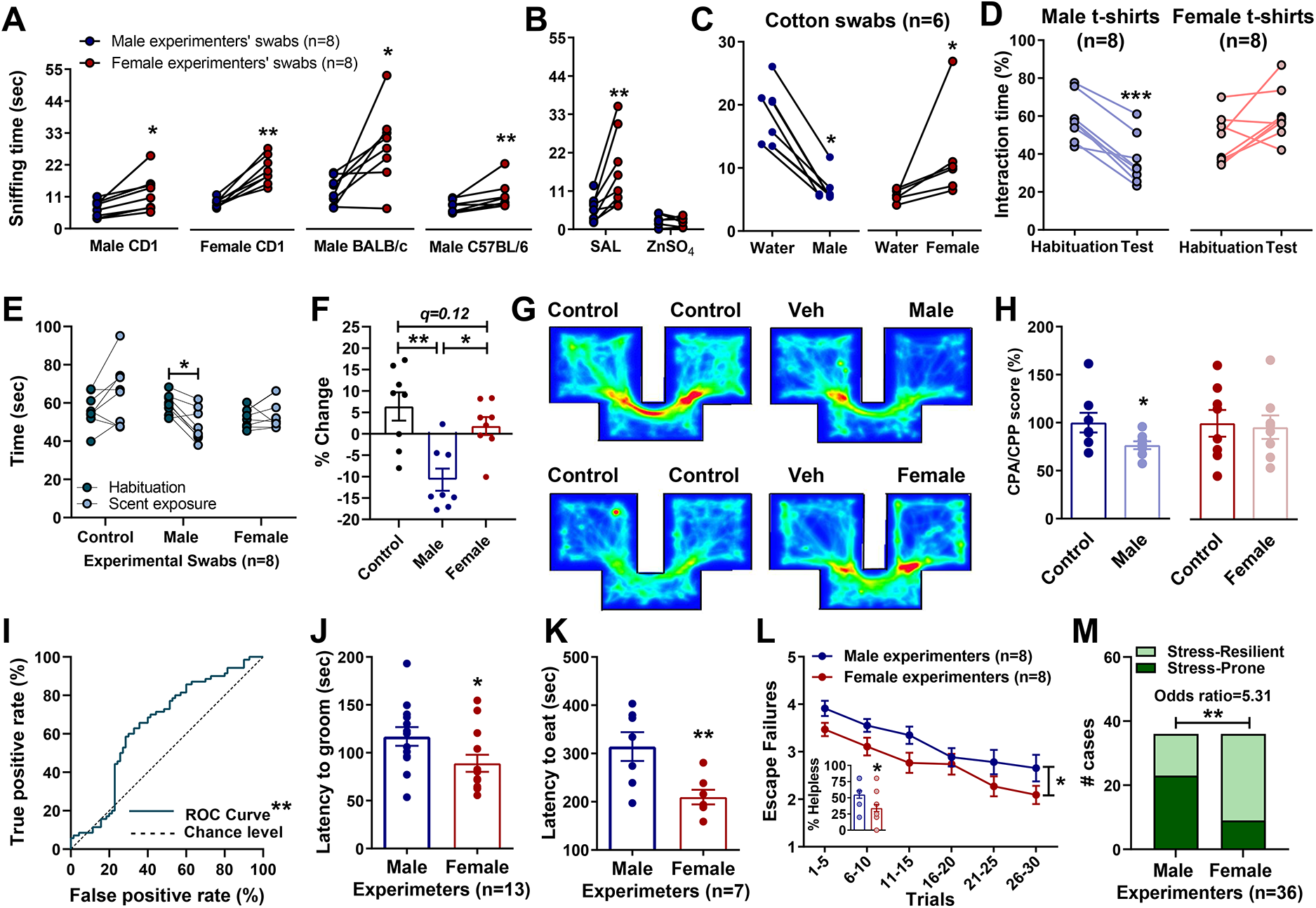
Mice manifest differential behavioural responses following exposure to male and female experimenter scent. Interaction time of **(A)** experimentally naïve and **(B)** anosmic mice with male *vs* female experimenter skin swabs. Time spent interacting with **(C)** male or female experimenters skin swabs *vs* control (water) swabs and **(D)** test *vs* habituation in the real-time place preference with experimenter and control t-shirts. **(E)** Time spent in each arm of a Y-maze with male, female and water scented swabs and **(F)** percent change from habituation. **(G)** Representative heat maps of **(H)** conditioned place preference/aversion of mice with male-paired and female-paired scents. **(I)** Receiver operating characteristic curve for prior experiments in this figure. **(J)** Latency to groom in the presence of male or female experimenters skin swabs. **(K)** Latency to eat in the novelty suppressed feeding test and **(L)** escape failures following inescapable shock training displayed by mice handled by male and female experimenters. (**M**) Contigency analysis of the number or stress resilient and stress prone mice. Data shown are mean ± S.E.M. * *p<*0.05; ** *p<*0.01; \*\*\**p<0.001.* Veh, vehicle; ZnSO_4_, zinc sulphate.

To determine whether sex-specific human scents affect stress-sensitive behavioural responses, we investigated the effect of exposure to male or female experimenters on forced-swimming test (FST) behaviours, where mice are placed into a cylinder containing water from which they cannot escape and their resilience to stop struggling is timed ^5^. Mice handled by male experimenters manifested significantly higher immobility time compared to those handled by female experimenters (Extended data Fig. 1A; pooled results from all FST experiments presented in this manuscript), indicative of a greater male experimenter-induced stress response. We also observed a higher latency to groom in the sucrose splash test, where the time to initiate grooming is measured after being sprayed with sucrose containg water, in the presence of male compared with female experimenter swabs (Fig. 1J) indicative of decreased self-care ^6^. Similarly, a greater latency to eat in the novelty-suppressed feeding test when the experiment was performed by male *vs* female experimenters (Fig. 1K), indicative of greater anxiety ^7^. Furthermore, increased escape deficits following exposure to inescapable foot shock training were observed when CD1 (Fig. 1L,M) or CFW Swiss-Webster (Extended data Fig. 1B,C) mice were handled by male *vs* female experimenters. Similarly, when a chronic social defeat stress (CSDS) paradigm, where mice are exposed once a day to a larger aggressive mouse for 10 days, was performed by a male experimenter, male C57BL/6J mice manifested significantly lower sucrose preference (indicative of anhedonia) compared to when the same experiment was concurrently performed by a female experimenter (Extended data Fig. 1D). In contrast, no significant difference was observed in the light-dark preference test in mice handled by male *vs* female experimenters (Extended data Fig. 1E).

### Sex of the human experimenter affects mouse responses to pharmacological treatments

Our identified effects of human experimenter sex on mouse behaviours may have implications beyond dimorphic behavioural responses. We assessed experimenter sex effects on responses to administration of (*R,S*)-ketamine (ketamine), a human clinically effective antidepressant at sub-anesthetic doses, which also exerts antidepressant-like actions in preclinical models ^8–10^. Under our routine laboratory testing conditions ^8, 11–13^, we found among five distinct pairs of experimenters that ketamine injected by male, but not female, experimenters decreased immobility time in male CD1 mice tested in the FST 1-hour (Fig. 2A). This experimenter sex-specific effect of ketamine was also observed 24-hours post-injection (Extended data Fig. 2A), in female CD1 mice (Extended data Fig. 2B), and in mice exposed to a pre-swim session 23 hours prior to the drug injection and 24 hours prior to re-exposure to the FST (Extended data Fig. 2C). In addition, ketamine administered by a male, but not a female experimenter, reversed CSDS-induced sucrose preference deficits (Fig. 2B; Days 1-3). After sucrose preference returned to normal levels, re-exposure to a brief (1 min) social defeat session resulted in a reinstated reduction in sucrose preference in both female and male experimenter-handled mice that had previously received vehicle (Fig. 2B; Days 9-12). However, mice that previously received ketamine treatment by a male, but not a female, experimenter demonstrated a protective phenotype against stress-reinstated sucrose preference deficits (Fig. 2B; Days 9-12). We also found when CD1 and CFW mice were administered ketamine by male, but not female experimenters, escape deficits induced by inescapable shock were significantly decreased 24 hours following treatment (Fig. 2C; Extended data Fig. 2D). Notably, replication of our ketamine findings was performed at an independent institution by eight male and eight female experimenters (Fig. 2D, Extended data Fig. 2E). In line with our observations, administration of ketamine decreased immobility time in the FST only when it was administered by male experimenters (Fig. 2D, Extended data Fig. 2E). The experimenter sex-induced differential antidepressant responses are neither due to altered pharmacokinetics of ketamine or its norketamine or (*2R,6R*; *2S,6S*) hydroxynorektamine (HNK) metabolites (Fig. 2E), nor to a shift in the dose response of ketamine (Extended data Fig. 2F) when administered by a female experimenter.

**Fig. 2:**
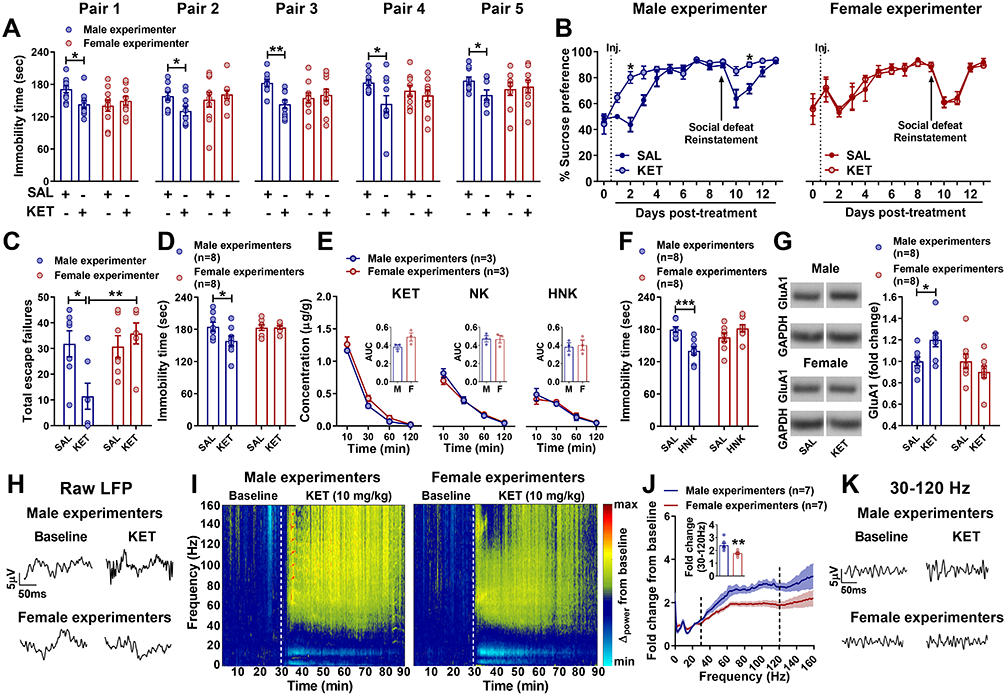
The sex of the human experimenter influences antidepressant and electroencephalographic responses to ketamine. **(A)** Immobility time in the forced-swim test (FST) performed 1-hour after dosing, **(B)** sucrose preference following social defeat, and **(C)** escape failures in the learned helplessness paradigm following either administration of ketamine (KET) or saline (SAL) by male *vs* female experimenter/s. **(D)** Immobility time in the FST 1-hour following either ketamine or saline injections by male or female epxerimenters at another institution. **(E)** Pharmacokinetic profile of KET, norketamine (NK), and (*2R,6R;2S,6S*)-hydroxynorketamine (HNK) following administration of ketamine (10 mg/kg, i.p.) by male or female experimenters. **(F)** Immobility time in the FST 1-hour following administration of (*2R,6R*)-HNK or saline by male or female experimenters. **(G)** Levels of GluA1 in ventral hippocampus. **(H)** Representative qEEG local field potentials (LFP), **(I)** spectrograms, **(J)** fold change from baseline, and **(K)** representative traces from 30-120 Hz following administration of ketamine by male or female experimenters. Data shown are mean ± S.E.M. * *p<*0.05; ** *p<*0.01

To determine whether experimenter sex-dependent behavioural responses to ketamine are specific for the antidepressant-relevant actions of the drug, and not its stimulant effects, we assessed hyperlocomotion following ketamine administration. Ketamine is an *N*-methyl-*D*-aspartate receptor (NMDAR) antagonist, which is responsible for many of the drugs side effects including stimulant actions ^14^. We observed an identical increase in NMDAR inhibition-dependent locomotor activity following ketamine administered by a male or female experimenter (Extended data Fig. 3A,B), suggesting sex of the experimenter-mediated differences in antidepressant-like responses are not directly related to NMDAR inhibition. Consistent with this, administration of the NMDAR antagonist, MK-801, by either a male or female experimenter decreased FST immobility time (Extended data Fig. 3C). (*2R,6R*)-HNK is a pharmacologically-active metabolite of ketamine that is currently in human clinical trials shares the antidepressant-like actions of ketamine ^8, 12, 14–20^, but has low potency for NMDAR inhibition ^16^. Administration of (*2R,6R*)-HNK by eight different male and female experimenters resulted in male experimenter-specific antidepressant-like behavioural responses in the FST, while no effect was observed when (*2R,6R*)-HNK was administered by female experimenters independent of dose (Fig. 2F; Extended data Fig. 3D,E). In contrast, administration of a classical tricyclic antidepressant, desipramine, resulted in similar antidepressant-like actions when administered by either a male or female experimenter (Extended data Fig. 3F). We also demonstrated that ketamine injected within a biosafety cabinet, to eliminate experimenter scent, did not result in antidepressant-like actions regardless of the sex of the experimenter (Extended data Fig. 4A). Altogether, these data support an interaction between exposure to male experimenters’ scent and ketamine’s antidepressant-like behavioural effects. In line with this, female experimenter-administered ketamine within a biosafety cabinet, but in the presence of male-worn t-shirts (an experimental manipulation to replace female experimenter scent with that of a male), led to a decrease in immobility time in the FST (Extended data Fig. 4B).

Increased hippocampal synaptic levels of the AMPA glutamate receptor GluA1 subunit have been demonstrated following ketamine administration, and AMPA receptor activity is required for ketamine’s antidepressant-relevant behavioural actions in rodents ^8–10^. Administration of ketamine by male, but not female experimenters, increased synaptic GluA1 AMPAR subunit levels in the ventral hippocampus of mice 24 hours post-treatment (Fig. 2G, Extended data Fig 4C). The ventral hippocampus can drive neuronal activity in the prefrontal cortex linked to ketamine’s antidepressant actions ^21^, which are mediated by AMPAR activity ^8^, leading us to assess prefrontal cortex oscillatory activity using quantitative electroencephalography (qEEG). As previously reported, ketamine increased qEEG power in the 30-120 Hz range (Fig. 2H-K). Notably, in line with evidence for an association between increased gamma power and ketamine’s antidepressant responses ^22^, we show that oscillations within the 30-120 Hz, power range occurred to a greater extent when ketamine was administered by male experimenters (Fig. 2H-K, Extended data Fig. 5A-F).

### Corticotropin-releasing factor interacts with ketamine to exert experimenter sex-specific physiological responses

The acute exposure of mice to male scent increases HPA axis activation evidenced by higher plasma corticosterone levels ^2^. However, inhibition of corticosterone synthesis by pre-treatment with metyrapone did not prevent the antidepressant-like actions of male-injected ketamine in CD1 mice (Extended data Fig. 6A). HPA-axis activation involves corticotropin-releasing factor (CRF) release, which precedes the corticosterone response ^23^. We thus investigated a possible interaction between experimenter scent, HPA axis activation, and antidepressant-like responses to ketamine. Intracerebroventricular (ICV) administration of CRF prior to administration of ketamine by a female experimenter within a biosafety cabinet resulted in decreased FST immobility 24-hr post administration in male and female CD1 mice (Fig. 3A) and decreased escape failures in the learned helplessness paradigm in ketamine- (Fig. 3B) and (*2R,6R*)- HNK-treated (Fig. 3C) mice. Consistent with these findings, peripheral administration of the brain-penetrant CRF1 receptor antagonist CP-154,526 blocked antidepressant-like actions of male experimenter-administered ketamine in the learned helplessness paradigm (Fig. 3D), FST (Fig. 3E), and sucrose preference test following CSDS (Fig. 3F). Furthermore, treatment with CP- 154,526 decreased aversion to male experimenters’ skin swabs (Fig. 3G,H) and abolished ketamine-induced increases in 30-120 Hz cortical oscillations (Fig 3I-L; Extended data Fig 7A-F).

**Fig. 3:**
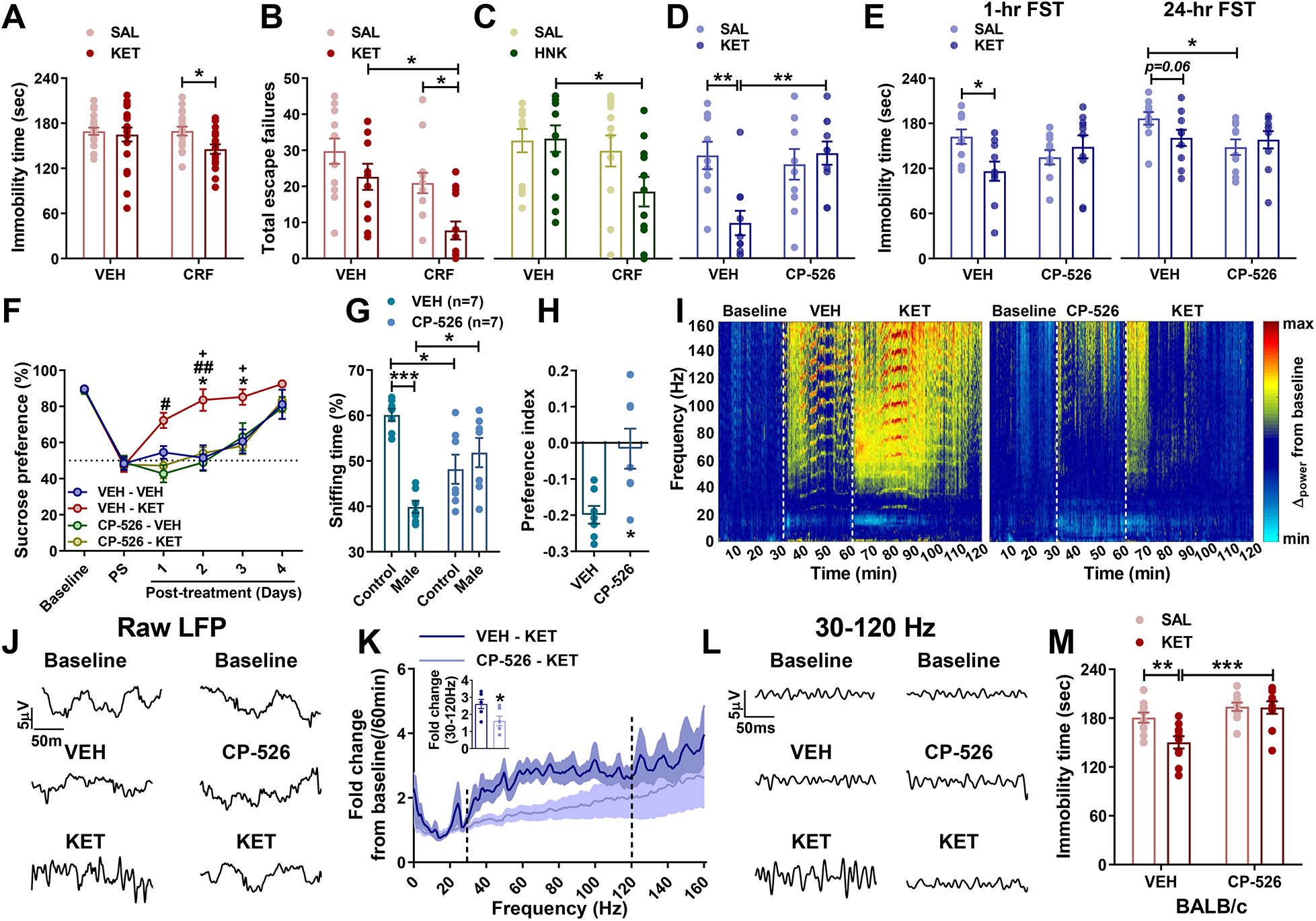
CRF mediates antidepressant responses to ketamine. Effects of intracerebroventricular administration of corticotropin-releasing factor (CRF) prior to ketamine (KET) in the **(A)** FST and **(B)** learned helplessness paradigm and **(C)** prior to (*2R,6R*)-hydroxynorketamine (HNK) administration in the learned helplessness paradigm by a female experimenter. **(D)** Escape failures, **(E)** immobility time, **(F)** chronic social defeat stress -induced sucrose preference deficits, and **(G)** skin swab preference following pretreatment with the CRF1 receptor antagonist CP-154,526 (CP-526) by a male experimenter. Representative **(I)** spectrograms and **(J)** traces from raw local field potentials (LFP), **(K)** fold change, and **(L)** representative 30-120 Hz traces derived from qEEG recordings from mice treated with CP- 526 prior to KET. **(M)** Effects of CP-526 administration prior to KET in BALB/c mice by a female experimenter. Data shown are mean ± S.E.M. * *p<*0.05; ** *p<*0.01; *** *p<*0.001.

We hypothesised the BALB/c strain of mice, which display higher baseline immobility time in the FST compared to CD1 and C57BL/6J mice, and greater sensitivity to the behavioural effects of CRF ^24, 25^, would manifest antidepressant-like responses to female-administered ketamine. Indeed, we observed that ketamine injected by both male and female experimenters decreased immobility time in the FST (Extended data Fig. 6B), and pre-treatment with CP-154,526 by a female experimenter blocked ketamine’s antidepressant-like effects (Fig. 3M) in BALB/c mice. Overall, these findings suggest that male scent may induce activation of CRF neurons, which results in a corticosterone synthesis-independent enhancement of stress responses; an effect critical for ketamine’s antidepressant-like actions.

To identify a mechanism underlying the interaction between CRF and antidepressant-related ketamine responses, we performed *in vitro* electrophysiological recordings of hippocampal CA1 field excitatory post-synaptic potentials (fEPSP) following stimulation of either the Schaffer collateral (SC) pathway or the projections from the entorhinal cortex (EC). Similar to our previously published findings ^8, 26^, we found that the biologically active ketamine metabolite (*2R,6R*)-HNK increases AMPAR-mediated fEPSPs following stimulation of the SC, but not the EC-CA1 projections (Fig 4 A,B; Extended data Fig. 8A,B). CRF has been reported to prime long-term potentiation in the hippocampus ^27, 28^, and thus, we hypothesised that CRF would enhance the fEPSP potentiation induced by (*2R,6R*)-HNK. While CRF did not further enhance SC-CA1 fEPSP responses induced by (*2R,6R*)-HNK (Extended data Fig. 8A,B; 9A-E), it induced a synergistic potentiation of (*2R,6R*)-HNK-non-responsive EC-CA1 synapses (Fig. 4C,D); an effect likely occuring due to an increase in presynaptic release probability evidenced by a decrease in paired-pulse ratio (PPR; Extended data Fig 10A-H). This EC-CA1 fEPSP potentiation (Fig 4E,F) and decrease in PPR (Extended data Fig 10 I-K) were blocked by prior bath application of CP-154,526. Use of CRF-Cre mice combined with a cre-dependent retrograde fluorescent label (AAV pCAG-FLEX-tdTomato-WPRE) injected into the vental CA1 identified CRF-expressing neurons projecting from the EC to CA1 (Fig 4G). We confirmed this finding using wildtype mice that received retrograde conjugated cholera toxin (CTb) injected to the ventral CA1 followed by RNAscope for CRF mRNA transcripts in the EC to test for colocalization of the CTb and CRF. We found that there is a ∼10-18% colocalization with no great differences between medial and lateral EC or between anterior and posterion EC (Extended data Fig, 11 A-H). We performed whole-cell electrophysiology in CRF-cre mice that received an injection of cre-sensitive channelrhodopsin-2 (ChR2) expressing virus to the EC (Fig 4H). Postsynaptic activation by blue light of the EC CRF projection terminals in CA1 was blocked by NMDA and AMPA receptor antagonists, revealing that this projection is excitatory (Fig 4I), which is in line with our fEPSP data demonstrating that the synaptic potentiation induced by CRF + (*2R,6R*)-HNK was blocked by DNQX (Fig 4C, E). We thus hypothesized that activation of the EC to CA1 CRF projection is nececarry for the antidepressant effects of ketamine. To test this, we optogenetically stimulated the CRF teminals of the EC to CA1 circuit in YFP or ChR2-injected mice for 5 min, immediadetly followed by administration of ketamine within the BSC. The combination of the blue light activation and ketamine administration resulted in decreased immobility time (Fig 4J) demonstrating that activation of this projection interacts with ketamine to exert its antidepressant-like effects.

**Fig. 4:**
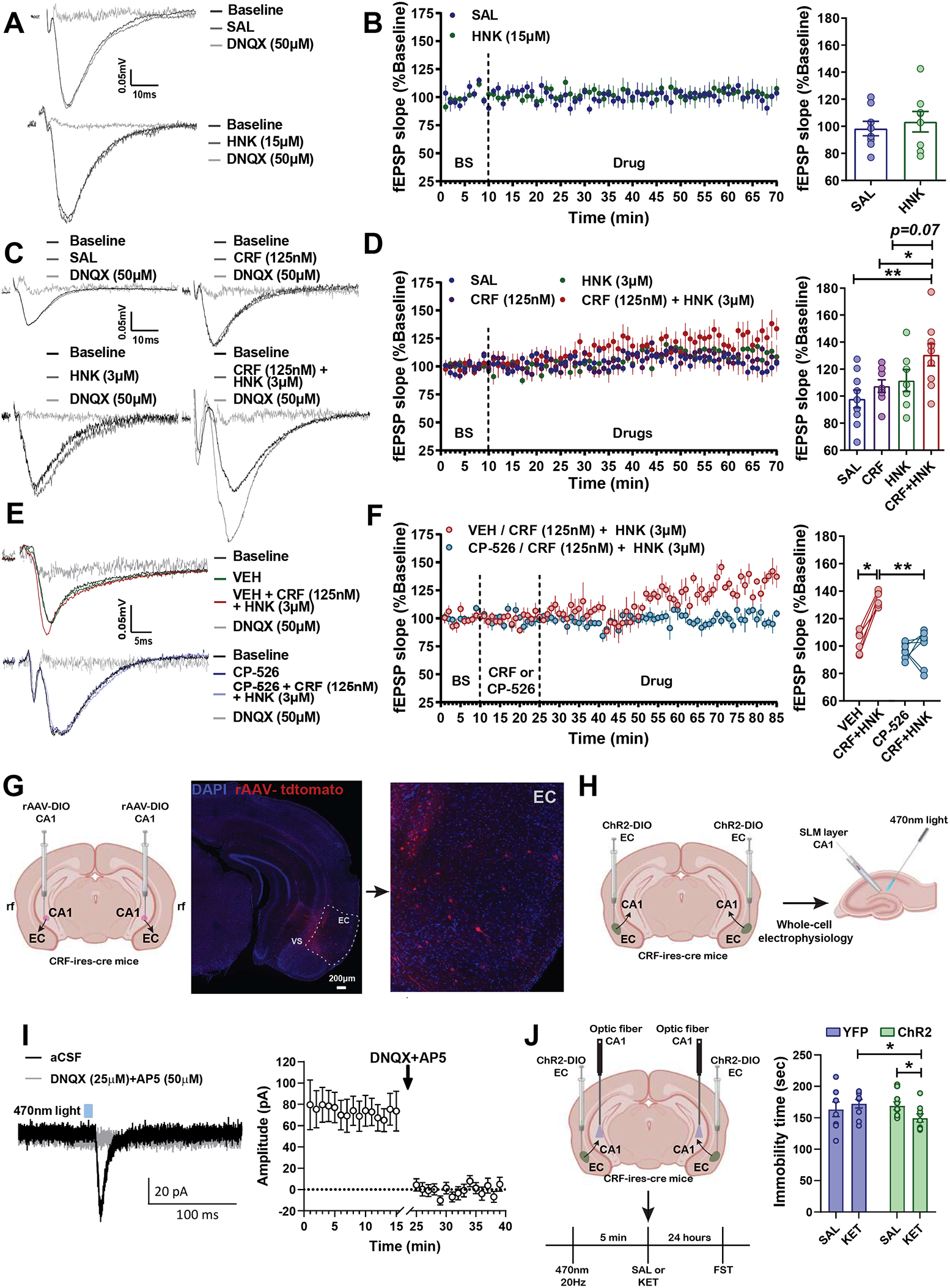
CRF mediates electophysiological responses to the (*2R,6R*)-hydroxynorketamine (HNK) ketamine metabolite. Representative traces and quantification of field excitatory postsynaptic potential (fEPSP) slopes following stimulation of the entorhinal cortex-hippocampal CA1 (EC-CA1) pathway following wash-in of **(A,B)** saline (SAL) and (*2R,6R*)-hydroxnorketamine (HNK) or in a separate experiment wash-in of **(C, D)** saline, corticotropin-releasing factor (CRF), (*2R,6R*)-HNK or CRF + (*2R,6R*)-HNK. **(E)** Representative traces and **(F)** quantification of fEPSPs slopes from the EC-CA1 pathway during pre-wash-in with the CRF1 antagonist CP-154,526 and wash-in with CRF + (*2R,6R*)-HNK. **(G)** Representative images from injection of retrograde AAV virus (tdTomaro) in CRF-cre mice to the CA1 area and labelled projections in the entorhinal cortex. **(H)** Experiment schematic, **(I)** representative traces and quantification of optogenetic stimulation to the CRF EC-CA1 circuit and whole cell recordings from postsynaptic cells. **(J)** Experiment Schematic and analysis of forced-swim test performance following the optogenetic stimulation of the CRF EC-CA1 circuit prior to ketamine administration within a biosafety cabinet. Data shown are mean ± S.E.M. * *p<*0.05; ** *p<*0.01; VS, ventral subiculum

Since EC is a region important in odour processing and discrimination ^29^, it led us to postulate that CRF neurons projecting from the EC to the hippocampus are activated in response to male scent, which in turn synergistically acts with (*2R,6R*)-HNK and ketamine to induce synaptic potentiation of that pathway. Indeed, we observed an increase in the number of CRF transcripts and CRF^+^/Fos^+^ co-labeled cells, but no difference in total Fos trancripts in the EC following exposure to male skin swabs (Fig 5A-D), suggesting that male scent induces activation of CRF neurons in the EC. To further investigate the involvement of this projection to the male experimenter aversion that we observed, we performed a real-time place preference experiment. ChR2 and YFP-injected mice received a light stimulation to the EC CRF terminals in CA1 when they were within the assigned compartment of the two compartment arena (Fig 5E). Activation of the CRF projection from EC to CA1 induced real-time aversion (Fig 5F), similar to the aversion observed in response to the scent of male experimenters (Fig 1E-F).

**Figure 5:**
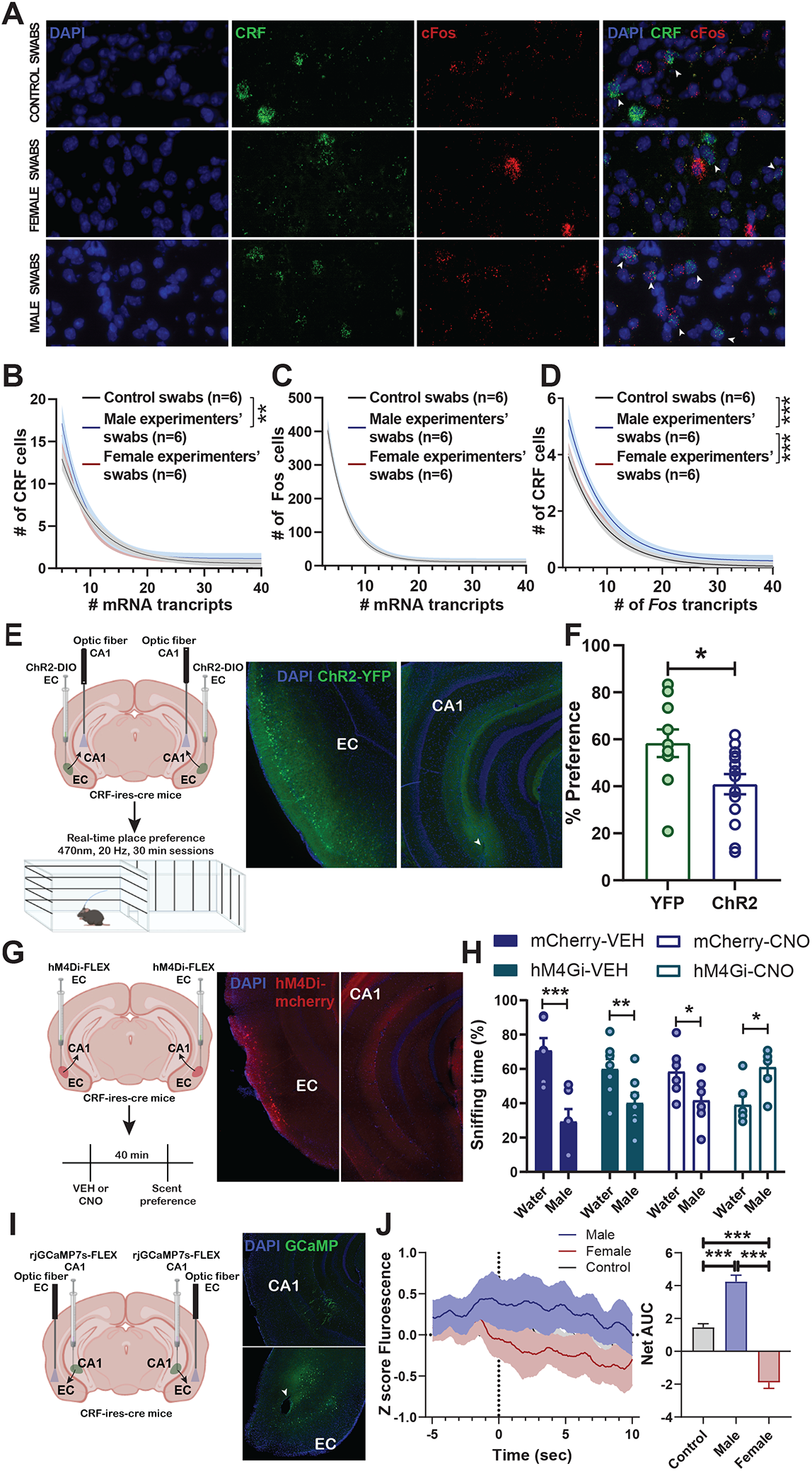
CRF positive EC cells mediate aversion to male experimenters’ scent. **(A)** Representative images and **(B-D)** quantification of CRF^+^/Fos^+^ cells in entorhinal cortex following exposure to experimenter scents. **(E)** Schematic of the experiment, represtative images **(F)** and quantification of the real time place preference during optogenetic stimulation of the CRF entorhinal cortex-hippocampal CA1 (EC-CA1) pathway. **(G)** Experiment schematic, represtative images and **(H)** quantification of the preference to the male experimenters’ scent compared to control follong chemogenetic inhibition of the CRF EC-CA1 pathway. **(I)** Experiment schematic and representative images and **(K)** quantification of the calcium transients of the soma of CRF EC-CA1 cells during a choice between male, female experimenter or water swabs using Fiber Photometry. Data for B, C and D are one-phase exponential decay curves representing the mean ± 95%. Data for E-K are shown as mean ± S.E.M for A-F. * *p<*0.05; ** *p<*0.01; *** *p<*0.001.

We next addressed whether inhibition of the activity of this projection, reversed the aversion to the male scent swabs (Fig 5G). This was accomplished by inhibition of the CRF projection form EC to CA1 through injection of cre-sensitive inhibitory DREADDs (hM4Gi-DIO-mCherry) into the EC of CRF-ires-cre mice and administration of CNO prior to exposure to the experimenter scent. We found that inhibition of the CRF EC neurons results in the reversal of male-scent aversion (Fig 5G,H) indicating that activity of CRF positive EC neurons are neccesary for adverse response to male scents (Fig 5G,H) suggesting that activity or EC-CA1 neurons will be associated with male scent exposure. In order to test this, we recorded calcium transients from the CRF EC to CA1 neurons using *in vivo* fiber photometry. CRF-ires-cre mice received an injection of a retrograde cre-sensitive jGCaMP7s bilaterally to the CA1. We then recorded from the CRF soma in the EC during exposure to male, female and control scent in a Y-maze as in Fig 1 E,F. CRF-cre mice were placed in a Y-maze equipment, each arm containing a male, female, or control swab while simultaneously recording neuronal activity (Fig 5I). We identified an increase in the activity of these neurons when mice were in the male scent arm compared to when they were within arms contains swabs exposed to either female and control scents (Fig 5J). Altogether these data indicate that the CRF EC to CA1 circuit is responsive to male scent.

Overall, we found that exposure to male or female experimenters’ scents induces differential baseline and stress-related behavioural responses, and alters responses to the antidepressant ketamine, in mice. Our findings are in line with earlier findings of Sorge et al demonstrating that exposure to the scent of male experimenters results in increased corticosterone, induction of stress-induced analgesia and an increase in anxiety-like behaviors ^2^. Our data further identify that exposure to male scent is aversive involving CRF expressing neurons that project from the EC to CA1. This specific brain circuit underlies the behavioral differences observed in response to male vs female experimenters.

Moreover, we find that the effects of experimenter sex not only affect baseline behaviors but also can impact responses to pharmacological treatments. This finding allowed for dissection of novel molecular mechanisms underlying ketamine’s pharmacological actions demonstrating exposure to male scent prior to ketamine administration activates the CRF system in the EC-CA1 pathway, which is necessary for ketamine’s antidepressant-relevant actions both *in vivo* and *in vitro*, and suggests a novel treatment approach for mood disorders involving combined treatment with ketamine/(*2R,6R*)-HNK and CRF agonists. Specifically we show that CRF activation of the EC to CA1 circuit is essential for the antidepressant effects of ketamine. This is likely via a cAMP-dependent mechanism relying on the Gs coupled CRF1 receptor, converging with our prior work implicating inhibition of Gi coupled mGluR2 ^12^, with effects of (*2R,6R*)-HNK to increase the probability of glutamate release identified here and previously published by our group ^26^.

Our findings highlight the relevance of the sex of the human experimenter on experimental outcomes, which may affect replicability, and the interpretation of results within and between laboratories. In our case male experimenter triggered stress responses resulted in an expected outcome (antidepressant-like effects of ketamine); however, this may be reversed for other studies. For example, it was previously observed that exposure to male experimenters resulted in stress-induced analgesia, which dampened the pain response in mice and rats ^2^. Understanding this general sexually dimorphic phenomenon, through its specific and quantitative contribution to different experimental outcomes, may lead not only to reduced heterogeneity between studies, but also increased capability to uncover novel biological mechanisms. Many other factors may affect behavioral results or contribute to the effects of experimenter sex (or lack thereof) that we observed such as ventilated *versus* open-air cages, whether mice were bred in a facility or shipped from commercial suppliers, the overall stress exposure within the facility, phase of day-night cycle, unique handling procedures, the strain and substrain of the mice used, stress level of experimenters, diet of experimenters (carnivore versus herbivore), or hormonal status among others.

## Materials and Methods

### Experimental design

All experiments were performed in a randomised manner (simple and stratified methods) by drug treatments and experimenter sex. Separate cohorts of mice were used for each experiment. Analysis of all experiments were performed blind to group assignments. Experimenters performing the injections and the behavioural paradigms were blind to the treatment groups. Experimenters were advised not to wear any perfumes or any perfumed cosmetics on the day of the experiments. All handling of mice and injections for behavioural experiments were performed in the animal vivarium. Mice were only handled by the assigned experimenters during the experiments. Experiments were determined not to require internal review board (IRB) approval with the condition that we neither collected nor associated any data from human experimenters other than their sex. All mice were weighed by the assigned experimenter the day prior to behavioural testing. No entrance was permitted during this time to any other person except for the experimenters. Injections by the male and female experimenters alternated every 30 min to ensure that the room was free from prior human experimenter scents. Independent cohorts of mice were used for all experiments. For experiments in Figure 1 and Figure 2D,F,G, H-K, Fig 3G-H and Fig 5A-D there were multiple male and female experimenters, each experimenter performed experiments with multiple mice (see *Supplementary Table 1* for exact numbers of experimenters and mice for each experiment) and the average outcome score of the mice from each experimenter was calculated (i.e. biological replicates) and used for statistical analyses. The numbers of experimenters and mice are provided in *Supplementary Table* 1. Experiments requiring a biosafety cabinet (BSC) were performed in a class II type A2 cabinet equipped with a HEPA filter. The supply filter was a HEPEX Seal with 99.99% effectiveness on 0.3 micron sized particles. The exhaust filter was a Neoprene, Springloaded filter with 99.99% effectiveness on 0.3 micron sized particles. The personal protective equipment of experimenters injecting under the BSC included single use labcoat on top of the regular clean labcoat and high-sleeve gloves that were worn above the labcoat’s sleeves.

### Animals

Mice (8 weeks old at arrival) were housed 4-5 per cage under a 12 h light–dark cycle (lights on at 7:00 am). Food and water were available *ad libitum*. Mice were acclimated to the animal vivarium for at least 7 days and experiments were performed in 9-12 weeks old animals. CD1, CFW and BALB/c mice were obtained from Charles River Laboratories (Raleigh, NC, USA), while C57BL/6J were obtained from a University of Maryland Baltimore Veterinary Resources mouse colony. CRF-ires-cre mice breeding pairs were kindly donated by Dr. Dennis Sparta (University of Maryland, Baltimore) and initially purchased from Jackson laboratories. Wild-type, heterozygous and homozygous CRF-ires-cre were produced in-house by breeding heterozygous males and females or by breeiding male and female homozygous CRF-ires-cre knockout mice with wildtype mice. Tail samples from CRF-ires-cre were obtained prior to weaning and genotyped by TransnetYX, Inc. (Cordova, TN, USA). All experimental procedures were approved by the University of Maryland, Baltimore or Yale University Animal Care and Use Committees and were conducted in full accordance with the National Institutes of Health Guide for the Care and Use of Laboratory Animals.

### Drugs

(*R,S*)-ketamine, desipramine, (+)-MK-801, CRF (h/r) (Sigma-Aldrich, St. Louis, MO, USA), (*2R,6R*)-HNK (National Center for Advancing Translational Science, Bethesda, MD, USA) were dissolved in 0.9% sterile saline. CP-154,526 (Abcam, Cambridge, UK) was dissolved in 20% ethanol and briefly heated at 60° C. Metyrapone (Cayman Chemica, Ann Arbor, MI, USA) was dissolved in 5% Tween 20. All drugs were administered intraperitoneally (i.p.) except CRF, which it was administered intracerebroventricularly (ICV) at a volume of 1 μg in 4 μl. Injection volume for all drugs was 7.5 ml/kg except CP-154,526 which was 1.5 ml/kg.

Drugs were administered at the following doses unless otherwise noted in the figures for the dose response experiments: (*R,S*)-ketamine (10 mg/kg), desipramine (20 mg/kg), (+)-MK-801 (0.03 mg/kg), CRF (h/r) (1 μg in 4 μl), (*2R,6R*)-HNK (10 mg/kg), CP-154,526 (30 mg/kg) and metyrapone (30, 50 and 70 mg/kg).

#### Surgical procedures

In all surgical procedures, mice were anesthetized with isoflurane at 3.5% and maintained at 2– 2.5% throughout the surgery. Analgesia was provided to the mice in the form of carprofen (5 mg/kg, s.c.; Norbrook laboratories, Newry, UK) prior the start of surgery and after the surgery once per day for three days.

### Male/female experimenter scent sniffing preference test

Clean cotton tipped applicators (VWR, Radnor, PA, USA) were dipped in distilled water and skin swabs were taken from *cubital fossa*, inner wrist, and mastoid, and immediately placed in separate air-tight bags by each experimenter using gloves. Applicators were used within 2 hours of collection. Two applicators were used per experimenter. Each scent-containing applicator (swab) was used for two mice. This experiment was performed in clean cages placed on air ventilated racks to avoid exposure of mice to other scents. First, mice were individually placed into clean cages for 10 min. Then, mice were habituated to a clean cotton tipped applicator for 30 min. Following habituation, the male- or female-experimenter-scented swabs were simultaneously introduced in the cage for 3 min. The time mice spent sniffing each swab was measured by an experimenter blind to the groups. The cages were only opened within the BSC when necessary for the experiment.

### Induction of anosmia

Zinc sulphate (ZnSO_4_) was used for the induction of anosmia. Mice were lightly anesthetised with isoflurane under the BSC and 10 μl of 5% ZnSO_4_ or saline was administered intranasally. Intranasal administration occurred one hour apart between each nostril. Behavioural testing occurred on the following day. Anosmia was confirmed by a food finding task immediately after the male/female experimenter scent sniffing test. The food provided was chocolate flavor sucrose pellets (Bioserv, NJ, USA) and placed in the front corner of each cage. All of the ZnSO_4_-treated mice were identified as anosmic (Mean latency ± S.E.M: SAL= 18.44 sec ± 14.69; ZnSO_4_ all mice reached the maximum testing time of 300 sec without finding/eating the food provided).

### Human experimenter *vs* control scent sniffing preference test

Clean cotton tipped applicators (VWR, Radnor, PA, USA) were dipped in distilled water and skin swabs were taken from cubital fossa, inner wrist and mastoid and immediately placed in an air-tight bag. Two applicators were used per experimenter. The control cotton tipped applicators were dipped in distilled water within the BSC. Each scent/control containing applicator was used for two mice. This experiment was performed in clean cages placed on air ventilated racks to avoid exposure of mice to other scents. First, mice were singly housed in a clean cage for 10 min. Then mice were habituated to a clean cotton tipped applicator for 30 min. Following habituation, the cotton applicators containing the human male or female and control scent were introduced in the cage for 3 min. The time mice spent sniffing each applicator was measured by an experimenter blind to the groups. The cages were only opened under the BSC when necessary for the experiment.

For the chemogenetic manipulation, CRF-ires-cre mice received an injection of 300nl of AAV5-hSyn-DIO-hM4D(Gi)-mCherry or AAV5-hSyn-DIO-mCherry (Addgene) into the EC (AP: -2.8, ML: ± 4.25, DV: - 5.17 from the top of the skull). Four weeks following the EC injection, mice received veh (7.5 ml/kg, ip) 40 minutes prior to scent introduction and one week later they were administered with CNO (3 mg/kg, ip). CNO was dissolved in 5% DMSO and injections were performed within a biosafety cabinet. Sessions were recorded with a video camera and analyzed by an experimenter blind to the experimental groups.

### Y-maze scent preference test

The Y-maze apparatus (Stoelting, IL, USA) consists of three identical arms (5 x 35 cm) joined in the middle, thus forming a ‘Y’ shape. Mice were placed in the center of the equipment and allowed to explore the apparatus for 8 minutes. Following habituation, swabs from male and female experimenters and water swabs were attached at the end of each arm. Mice were re-introduced to the arena, now including the swabs for 8 minutes All the experimental trials were recorded with an overhead videocamera and analyzed with TopScan (CleverSys) tracking software. The time spent in the area close to the swab, 75% of the arm, was calculated.

### Scent-induced real-time place preference

The real-time place preference apparatus consisted of a rectangular three-chambered box (40 cm length × 30 cm width × 35 cm height; Stoelting, IL) divided into three compartments (two equal-sized end-chambers and a middle chamber). A clear plexiglass perforated divider was placed in each end-chamber separating them into two compartments. The t-shirts were placed in the compartment (10 x 15 x 36 cm) located towards the end of the chamber away from the middle chamber so that the mice could not have direct physical interaction with the t-shirts. For this experiment we used t-shirs worn for 24 hours by the experimenters. The t-shirts were never worn by anyone else prior to this experiment. Experimenters were adviced to place the t-shirts in an air-tight bag in the morning of the experiment. During the real-time place preference, the mouse was initially placed in the apparatus without the t-shirts and allowed to freely explore for 2.5 min. Following that, the t-shirts were introduced, and the mouse was let to explore for another 2.5 min. The control and male- or female-worn t-shirts were counterbalanced with respect to chamber side during the experiment. A male experimenter performed the experiment with the male worn t-shirts and a female experimenter performed the experiment with the female worn t-shirts. The exprimenters were immediately leaving the testing room upon placement of the mice into the behavioural chamber. Each t-shirt was used for two mice and the average scores of those two mice was calculated for the final results. The real-time place preference experiment was performed under low-lighting conditions (30 lux) and sessions were recorded using overhead cameras. An experienced experimenter blind to the experimental groups scored the amount of time mice sniffed the experimenter-worn t-shirt *vs* the control t-shirt.

### Optogenetic induced Real-time place preference

A three-chambered box (40 cm length × 30 cm width × 35 cm height; Stoelting, IL) was used to measure the preference of mice to the light-paired compartment with the non-light paired compartment. One compartment was altered with white vertical stripes 4 cm apart on each wall and a smooth grey floor, while the other had white horizontal stripes 4 cm apart on the walls and a smooth grey floor. Allocation of the light-paired compartment was counterbalanced to avoid any side preferences. Heterozygous CRF-ires-cre mice received injections of 300nl of AAV5.EF1a- DIO-eYFP-WPRE-hGH or AAV5.EF1a-DIO-hChR2(H134R)-eYFP-WPRE-hGH; Penn Vector Core and Addgene) in the EC (AP: -2.8, ML: ± 4.25, DV: - 5.17 from the top of the skull) at a rate of 20nl/min using a 1µL Hamilton Syringe (Knurled Hub Needle, 25G) followed by implantation of the optic cannulas (Neurophotometrics, San Difego, CA) in the CA1 (AP: -2.3; ML: ±3.25; DV: -4.5 from top of the skull) three weeks after virus infusion. Ten to fifteen days following implantations ChR2 and YFP-injected mice were connected to optogenetic fiber cables (Doric Lenses Inc., Quebec, Canada) and placed in the middle compartment, and tracked by an overhead camera. During the 30 min testing period, mice that crossed into the allocated light-paired compartment received a 20Hz continuous stimulation (470nm, 200μm, 0,37NA, ∼2-3mW). Mouse crossings into each compartment were detected by a Bonsai script which communicated to the LED through through connected with a PulserPlus box (Prizmatix, Givat-Shmuel, Israel) for light delivery.

### Conditioned-place preference/aversion

The same three-chambered apparatus as the one described for the real-time place preference experiment (40 cm length × 30 cm width × 35 cm height; Stoelting, IL), was also used for the conditioned-place preference/aversion paradigms. Removable doors were used to separate the apparatus into two equal sized compartments. One compartment consisted of white vertical stripes 4 cm apart and a smooth black floor and the other compartment consisted of black walls and a perforated floor. The protocol consisted of a pre-conditioning phase, three conditioning sessions and a post-conditioning test. During the pre-conditioning day, mice were placed in the middle chamber and allowed to freely explore the apparatus for 10 min. The conditioning phase consisted of three days with two conditioning sessions per day. During these sessions, mice were confined to a single compartment and exposed to human experimenter or control scent; two cotton tipped applicators containing experimenter scents were secured on the walls, 15 cm above the floor using clear tape, of one of the compartments during the morning sessions and control (water) cotton applicators were placed in the opposite compartment during the afternoon sessions. Conditioning sessions lasted 10 min and the top of the apparatus was covered using black Plexiglass to keep the smell of the swabs in the chamber as much as possible. Female scent was paired with the least preferred compartment (defined during the pre-conditioning test) to assess for place preference and male scent with the most preferred compartment to assess for place aversion. Mice were exposed to different (same-sex) experimenter scent during each conditioning session. Collection of the cotton-tipped applicators containing human experimenter scent was performed as described earlier. During the post-conditioning test, mice were placed in the center chamber and allowed to freely explore the whole apparatus for 10 min. Time spent in each compartment was analyzed using TopScan video tracking software (CleverSys) during both the pre- and post-conditioning sessions for the last five min, as previously described (*1*).

### Sucrose splash test

Mice were placed in clean cages without bedding. Following 10 min of habituation in the cage, swabs (two cotton-tipped applicators) from either a male or female experimenter were attached on the inside of the cage top using clear tape. The mice were sprayed with a 10% sucrose solution on their dorsal coat surface and latency to groom was scored by an experimenter blind to the treatment groups. This experiment was perform under the BSC.

### Novelty-suppressed feeding test

Mice were singly housed and food-deprived overnight in freshly made home-cages. One regular chow diet pellet was placed on an inverted petri-dish platform in the center of an open-field arena (40 × 40 cm). On the testing day, male and female experimenters placed their assigned mice in the boxes and left the room. Sessions lasted for 15-min and were recorded by video cameras for analysis. The latency for mice to take a bite of food was scored by a trained experimenter blind to the experimental groups.

### Light/Dark box

Male or female experimenters placed mice in the illuminated compartment of the light/dark box (35 × 35 cm), facing the wall opposite to the dark compartment, and allowed to explore the whole apparatus for 10 min. The sessions were recorded using overhead video-cameras and the time spent in the illuminated and dark compartment was scored using TopScan v2.0 (CleverSys, Inc., Reston VA). Each experimenter was assigned a cage of two male CD1 mice.

### Footshock-induced escape deficits and reversal of escape deficits

#### Induction

The behavioural testing consisted of two phases, the inescapable shock training and screening for helplessness. Eight male and female experimenters were assigned with two cages of five CD1 mice (5 male and 5 female mice). For the inescapable shock training (Day 1), the animals were placed in one side of two-chambered shuttle boxes (Coulbourn Instruments, Whitehall, PA, USA), with the door between the chambers closed. Following a 5-min adaptation period, 120 inescapable foot- shocks (0.45 mA, 15-sec duration, randomised average inter-shock interval of 45 sec) were delivered. During the screening session (Day 2), mice were placed in one of the two chambers for a 5 min adaptation period. Following the adaptation period, a shock (0.45 mA) was delivered, with the door between the two chambers opening simultaneously. If the animal crossed over to the adjacent chamber the shock was terminated; whereas if the animal did not cross over, the shock was terminated after 3 sec. A total of 30 screening trials of escapable shocks were presented with an average of 30-sec delay between each trial. Mice were considered helpless if they had >5 escape failures during the last 10 screening shocks.

#### Reversal

For the escape deficits reversal experiments we used the same paradigm as described by Zanos *et al*., 2016. Briefly, the protocol consisted of four phases: inescapable shock training, screening, treatment and the test. The training (Day 1) and screening (Day 2) sessions were performed as described above for the induction of escape deficits. Mice that developed escape deficits (>5 escape failures during the last 10 screening shocks; helpless mice) were given an injection of either saline or ketamine on Day 3. CP-154,526 was administered 30-min prior to saline/ketamine and CRF was preceding saline/ketamine by a few sec. During the testing phase (Day 4), the animals were placed in the shuttle boxes and, after a 5-minute adaptation period, a 0.45-mA shock was delivered concomitantly with door opening for the first 5 trials, followed by a 2 sec delay for the next 40 trials. Similar to the screening session, crossing over to the sec chamber terminated the shock, while if the animal did not cross the shock was terminated after 24 sec. Each mouse received a total of 45 trials of escapable shock with an average of a 30 sec inter-trial intervals. The number of escape failures was automatically recorded (Graphic State v3.1; Coulbourn Instruments, Allentown, PA, USA).

All the phases of the footshock-induced escape deficits protocol, as well as the treatments were performed by the same experimenter. Slots by both male and female experimenters were run interchangeably with 30-min time gap between experimenters to ensure that the room and equipment was human experimenter scent-free.

### Forced-swim test (FST)

Mice were tested in the FST 1 hour or 24 hours post-injections. Pre-treatments with metyrapone and CP-154,526 occurred 30 min prior to ketamine/saline treatments under the BSC. CRF (ICV) was administered at the same time as ketamine under the BSC; ICV administration was performed as previously described ^13^. Due to the use of a very brief isoflurane anesthesia, CRF-treated mice were tested only in the FST 24 hours after drug adminsitration. For the optogenetic stimulation, heterozygous CRF-ires-cre mice received injections of 300 nl of AAV5.EF1a-DIO-eYFP-WPRE-hGH or AAV5.EF1a-DIO-hChR2(H134R)-eYFP-WPRE-hGH; Penn Vector Core and Addgene) to the EC (AP: -2.8, ML: ± 4.25, DV: - 5.17 from the top of the skull) at a rate of 20nl/min using a 1µL Hamilton Syringe (Knurled Hub Needle, 25G) followed by implantation of the optic cannulas (Neurophotometrics, San Difego, CA) in the CA1 (AP: -2.3; ML: ±3.25; DV: -4.5 from top of the skull) three weeks after virus infusion. Ten to fifteen days following implantations ChR2 and YFP-injected mice were connected to optogenetic fiber cables (Doric Lenses Inc., Quebec, Canada) YFP and ChR2 mice received a 20Hz stimulation of 470nm light ∼2-3mW (200μm, 0,37NA; Doric Lenses Inc., Quebec, Canada)for 5 minutes prior to the injection with ketamine. Mice were test in the FST 24-hours post-injection. The FST was performed as previously described (*3*). Briefly, mice were subjected to a 6-min FST session in clear Plexiglas cylinders filled to 15 cm with water (23 ± 1° C) under normal lighting conditions (800 lux). The FST was recorded using a digital video camera. The FST was always performed by a female experimenter who was not giving injections in that particular experiment. Immobility was defined as passive floating with no additional movements other than those necessary to stay afloat. Immobility time was scored for the last 4 min by a trained observer blind to the experimental groups.

In the 2-day FST experiment, mice were exposed to a 15-min pre-swim session on day 1. On day 2, mice were exposed to a 6-min swim session as described above. Ketamine was administered 1 hour prior to the testing session which was performed on day 2 and 23 hours after the pre-swim session (which was performed on day 1). Immobility time was scored for the last 4 min of the 6- min session on day 2 by a trained observer blind to the experimental groups.

In the FST experiment in which t-shirts were used, male experimenters were asked to wear the t-shirts for a 24 hour period, after which t-shirts were placed in an air-tight bag until use. Contol t-shirts were not worn by anyone. T-shirts were placed under the biosafety cabinet and mice were scruffed for injections on the male-worn t-shirts by a female experimenter (so that mice could only smell the t-shirt and not the person injecting). One-hour post-injection, mice were tested in the FST.

### Chronic social defeat stress

Male C57BL/6J mice underwent a 10-day chronic social defeat stress paradigm, as previously described ^8, 12^, with minor modifications. Briefly, experimental mice were introduced to the home cage of a resident aggressive retired CD1 breeder for 10 min. Following this physical attack phase, mice were transferred and housed in the opposite side of the resident’s cage divided by a Plexiglas perforated divider, in order to maintain sensory contact, for the remainder of the day. This process was repeated for 10 days. Experimental mice were introduced to a novel aggressive CD1 mouse each day. After the last defeat session on Day 10, the C57BL/6J mice were singly-housed in clean cages and returned to the animal vivarium. From day 10 to 12 mice were tested for the development of anhedonia by assessing their preference to sucrose. In particular, two pre-weighed bottles, one containing water and the other a 1% (w/v) sucrose solution, were introduced on the experimental C57BL/6J mice cage. Water and sucrose consumption by weight was measured every day. Mice that had sucrose preference lower than 60% were given a single injection of either saline (7.5 ml/kg) or ketamine (20 mg/kg) on day 12. Sucrose and water consumption was measured for an additional 9 days, at which time all mice recovered their preference. Then, on Day 21, experimental C57BL/6J mice were placed in the home-cage of a novel aggressive CD1 for 1 min to assess for reinstatement of sucrose preference deficits and possible prophylactive actions of prior ketamine administration. Sucrose consumption was measured daily for another four days.

The chronic social defeat stress paradigm was run in parallel by a male and female experimenter in two separate behavioural rooms, adjacent to each other using the same lighting and temperature/humidity conditions. Following the termination of the social defeat and the return of the mice to the animal vivarium for sucrose preference testing, the initial male and female experimenters were the only ones to open the cages and measure the sucrose and water consumption. Sucrose preference measurements were performed 30 min apart between each experimenter. The social defeat reinstatement phase was performed in two separate rooms by the same initial experimenters.

For testing the role of the CRF1 receptor on male-injected ketamine reversal of CSDS-induced sucrose preference deficits, only mice that had baseline preference higher than 80% were used. The CSDS procedure was performed as described above and the mice were exposed only to the male experimenter running the experiment. CP-154,526 was administered 30 min prior to ketamine/saline administration.

### Open-field test

Mice were placed into individual open-field arenas (50 x 50 x 38 cm; San Diego Instruments) for a 30-min habituation period and then received a saline injection by a male or female experimenter and assessed for locomotor activity for further 30 min. Then, mice received a ketamine injection (10 mg/kg; i.p) by a male or female experimenter during consecutive sessions and were assessed for locomotor activity for 60 min. Distance travelled and time spent in the center of the open-field arena was analyzed using TopScan v2.0 (CleverSys, Inc.).

### Western blots

Male CD1 mice received a ketamine (10 mg/kg; i.p.) or vehicle injection by either male or female experimenters and were euthanised via cervical dislocation 24 hours post-treatment. The ventral hippocampi were dissected and homogenized in Syn-PER Reagent (ThermoFisher Scientific, Waltham, MA, USA; 87793) with 1× protease and phosphatase inhibitor cocktail (ThermoFisher Scientific, Waltham, MA, USA; 78440) to purify synaptoneurosomes. The homogenate was centrifuged for 10 min at 1,200g at 4° C. The supernatant was centrifuged at 15,000g for 20 min at 4° C. After centrifugation, the pellet (synaptosomal fraction) was re-suspended and sonicated in N-PER Neuronal Protein Extraction Reagent (ThermoFisher Scientific, Waltham, MA, USA; 87792). Protein concentration was determined via the BCA protein assay kit (ThermoFisher Scientific, Waltham, MA, USA; 23227). Equal amount of proteins (10–40 μg as optimal for each antibody) for each sample was loaded into NuPage 4–12% Bis-Tris gel for electrophoresis. Gel transfer was performed with the TransBlot Turbo Transfer System (Bio-Rad, Hercules, CA, USA) Nitrocellulose membranes with transferred proteins were blocked with 5% milk in TBST (TBS plus 0.1% Tween-20) for 1 hour and incubated with primary antibodies overnight at 4° C. The following primary antibodies were used: GluA1 (Cell Signaling Technology, Danvers, MA, USA; 2983) and GAPDH (Abcam, Cambridge, UK; ab8245). The next day, blots were washed three times in TBST and incubated with horseradish peroxidase conjugated anti-mouse or anti-rabbit secary antibody (1:5,000 to 1:10,000) for 1 hour. After three final washes with TBST, bands were detected using enhanced chemiluminescence (ECL) with the Syngene Imaging System (G:Box ChemiXX9). After imaging, the blots were incubated in stripping buffer (ThermoFisher Scientific, Waltham, MA, USA; 46430) for 10–15 min at room temperature followed by three washes with TBST. The stripped blots were incubated in blocking solution for 1 hour and incubated with the primary antibody directed against total levels of the respective protein or GAPDH for loading control. Densitometric analysis of phospho- and total immunoreactive bands for each protein was conducted using Syngene’s GenTools software. Protein levels were normalised to GAPDH. Fold change was calculated by normalisation to saline-treated control group.

### Tissue distribution and clearance measurements of ketamine and metabolites

Male CD1 mice received a ketamine (10 mg/kg; i.p.) injection by either a male or female experimenter and were decapitated at 10, 30, 60 or 240 min post-injection following a 2-min exposure to 3% isoflurane. Trunk blood was collected in EDTA-containing tubes and centrifuged at 5938g for 6 min at 4°C. Plasma was collected and stored at −80°C until analysis. Whole brains were simultaneously collected, immediately frozen on dry ice, and stored at −80°C until use. The euthanasia and sample collection was performed by the same experimenter that performed the injections. Three separate cohorts were performed with three different pairs of experimenters.

Achiral liquid chromatography–tandem mass spectrometry was used to determine the concentration of ketamine and its metabolites in plasma and brain tissue as previously described (*27*). Brains were suspended in 990 μl of water:methanol (3:2, v/v) with the addition of D_4_- ketamine (10 μl of 10 μg/ml). The subsequent mixture was homogenized on ice with a polytron homogenizer and centrifuged at 21000 × *g* for 30 min. The supernatant was processed using 1-ml Oasis HLB solid-phase extraction cartridges (Waters Corp., Waltham, MA), which were previously pre-conditioned with 1 ml methanol, followed by 1 ml water and then 1 ml ammonium acetate (10 mM, pH 9.5). Following the addition of the supernatants to the cartridges 1 ml water was added, and the compounds were eluted with 1 ml methanol. The eluent was transferred to an autosampler vial for analysis. Quality control standards were prepared at 200, 500 and 1000 ng/ml.

For plasma samples, the calibration standards for (*R,S*)-ketamine, (*R,S*)-norketamine and (*2R,6R*;*2S,6S*)-HNK ranged from 5,000 to 19.53 ng/ml. The quantification of (*R,S*)-ketamine, (*R,S*)-norketamine, and HNK was accomplished by calculating area ratios using D_4_-ketamine (10 μl of 10 μg/ml solution) as the internal standard. AUC was calculated using Graphpad Prism v8.21.

### Quantitative Electroencephalogram

Quantitative Electroencephalogram (qEEG) experiments were performed according to previously published methods ^8, 12^, with minor modifications. Mice were anesthetised with isoflurane at 3.5% and maintained at 2–2.5% throughout the surgery. Analgesia was provided to the mice (carprofen, 5 mg/kg, s.c.) prior the start of surgery. An ETA-F10 radio-telemetric transmitter (Data Sciences International) was placed subcutaneously and its leads implanted over the dura above the frontal cortex (1.7 mm anterior to bregma) and the cerebellum (6.4 mm posterior to bregma). Surgeries were performed by either female or male experimenters. Animals were allowed to recover for 7 days before recordings. Mice were singlyhoused and placed in the behavioural room immediately after the surgeries. qEEGs were recorded using the Dataquest A.R.T. acquisition system and Ponemah version 6.32 acquisition systems (Data Sciences International) with frontal qEEG recordings referenced to the cerebellum.

Male and female experimenters injected ketamine (10 mg/kg; i.p.) with a 30 min gap time between them. Baseline (pre-injection) recordings were analyzed for 30 min prior to the ketamine injections for each experimenter. A further 60 min analysis was performed following ketamine injection. qEEGs were analyzed using Neuroscore version 3.2.9297-1 (DSI, MN, USA). An automated analysis protocol was implemented to mark occurrences of invalid signal data/artifacts. These artifacts were defined using an amplitude detector set at greater than or equal to an absolute threshold of 0.001 mV, with a minimum duration of 1E-05 sec, maximum duration of 10 sec, join interval of 1 sec, prepend duration of 0.01 sec, and append duration of 0 sec. qEEG signals were exported as periodograms using the following parameters: values to 6 decimal places using 10-sec epochs, Hamming windows, a Fast Fourier Transform order of 10, 50% overlap, and excluding epochs overlapping invalid data markers. These parameters yielded periodograms with frequency bins of 0.98 Hz. Oscillation power in each bandwidth (delta = 1–3 Hz; theta = 4–7 Hz; alpha = 8– 12 Hz; beta = 13–29 Hz; gamma = 30–100; HFO: 100-160 Hz) was calculated in 10-min bins.

Spectrograms expressed in log scales and indicated as heat maps were created in MATLAB 2018a by generating average periodograms within treatment groups and normalizing over mean treatment group baseline values for each frequency bin. Data for line graphs displaying average power changes across frequencies were calculated in MATLAB by dividing power values averaged across 30 min post-treatment by power values averaged across the baseline for each animal in each frequency bin, and graphs with S.E.M. were generated using GraphPad Prism version 8.3.

### Field excitatory postsynaptic potentials

Mice were euthanised by a brief exposure to isoflurane, followed by decapitation. Removal of the mouse brains and extraction of the hippocampi were performed in ice-cold ACSF bubbled with 95% O2, 5% CO2. The ACSF contained (in mM): 124 NaCl, 3 KCl, 1.25 NaH2PO4, 1.5 MgCl2, 2.5 CaCl2, 26 NaHCO3 and 10 glucose. Hippocampal slices were cut at 400 μm using a vibratome and kept in a holding chamber at the interface of ACSF and humidified 95% O_2_, 5% CO_2_ for at least 1 h. For the fEPSPs, slices were transferred to a submersion-type recording chamber and perfused with ACSF (0.5–2 ml/min; room temperature). Concentric bipolar tungsten electrodes were placed in stratum radiatum to stimulate the Schaffer collateral (SC) afferents and in the in stratum lacunosum-moleculare to stimulate entorhinal cortex (EC) afferents. Extracellular recording pipettes were filled with ACSF (3–5 MΩ) and placed in stratum radiatum for the SC and in stratum lacunosum-moleculare for the EC-CA1 pathway stimulation. Field potentials were evoked by monophasic stimulation (100 μs duration) at 0.1 Hz. Baseline stimulus strength was set to ∼40-50% maximum of field potential response to facilitate observing an increase in excitability and to avoid a possible ceiling effect. A stable baseline was recorded for at least 10 min. Vehicle, CRF (125nM) and (*2R,6R*)-HNK (3 or 15μM) were applied by perfusion over a period of 1-hr. For AMPAR-mediated responses, peak fEPSP slopes, measured over a window of 1–4 ms following the rising phase of the response, are reported as percentage change from baseline. DNQX (50 μM) was bath-applied at the end of the recording to ensure AMPA-mediated responses. Treatments were randomly assigned prior to the beginning of the recordings; however, the experimenter was not blind to the treatment groups during the experiment. Analysis of the fEPSPs was performed from an experimenter blind to the treatment groups. Slices which demonstrated paired-pulse ratio above 6 either before or after drug wash-in were deemed unhealthy and excluded from all analyses. The cases were the number of PPR data do not match the numbers of slices used during the potentiation experiments occurred because PPR protocol was mistakenly not performed. Experiments were performed and analysed using pCLAMP software (Molecular Devices, San Jose, CA, USA). Statistical analysis for the potentiation experiments was performed using the average of the last 5 minutes of the wash-in period. In the experiment were CP-154,526 was used, comparisons were performed between the average of the last 5-min of the pre-wash-in and the last 5-min of the wash-in period.

### Whole cell slice electrophysiology

Heterozygous CRF-ires-cre mice were used that previously received an injection of 300nl AAV5.EF1a-DIO-eYFP-WPRE-hGH or AAV5.EF1a-DIO-hChR2(H134R)-eYFP-WPRE-hGH; Penn Vector Core and Addgene) into the EC (AP: -2.8, ML: ± 4.25, DV: - 5.17 from the top of the skull) at a rate of 20nl/min using a 1µL Hamilton Syringe (Knurled Hub Needle, 25G). Five weeks following injections, mice were decapitated and their brains were rapidly removed and transferred to an oxygenated (95% O_2_, 5% CO_2_) ice-cold solution containing (in mM): 30 NaCl, 2.5 KCl, 1.25 NaH_2_PO_4_, 26 NaHCO_3_, 10 MgCl_2_, 0.5 CaCl_2_, 10 Glucose, 230 Sucrose. Coronal slices containing the hippocampus (250µm) were sectioned using a Leica VT1200S vibratome (Leica Biosystems) and transferred to a holding chamber filled with oxygenated solution containing (in mM): 109 NaCl, 4.5 KCl, 1.2 NaH_2_PO_4_, 35 NaHCO_3_, 1 MgCl_2_, 2.5 CaCl_2_, 11 Glucose, 20 HEPES, 0.4 Ascorbic acid. Slices were left at room temperature (RT) in the holding chamber at least 1 hour before the start of the experiments. After about 1 hour, slices were transferred to a small-volume (<0.5 mL) recording chamber that was mounted on a fixed-stage upright microscope (Nikon Eclipse E600FN) equipped with differential interference contrast (DIC) optics and immersed in oxygenated aCSF containing (in mM): 125 NaCl, 2.5 KCl, 1.25 NaH_2_PO_4_, 26 NaHCO_3_, 1.5 MgCl_2_, 2.4 CaCl_2_, 25 Glucose. (for all solution: pH 7.4, mOsm about 310). Experiments were performed at RT and the aCSF was flowing at 2 mL/min. Neuronal somata were visualized at high magnification (40x, 0.8 NA water-immersion objective; Nikon). Conventional tight-seal (>2 GΩ) whole cell patch-clamp recordings were acquired using Multiclamp 700B (Molecular Devices) amplifier. The signal was filtered at 1 kHz and digitized at 10 kHz using Axon Digidata 1550B (Molecular Devices). Recording pipettes (3.5-5 MΩ) were pulled with P-97 horizontal micropipette puller (Sutter Instruments) filled with internal solution containing (in mM): 100 CsCH_3_SO_3_, 50 CsCl, 10 HEPES, 2 MgCl_2_, 0.2 EGTA, 10 Phosphocreatine (Tris), 4 Mg-ATP, 0.3 Na-GTP. 1.5 mg/mL QX-314 was added to the internal solution on the day of the experiment. (pH was adjusted to 7.2-7.3 with CsOH, mOsm 285-290). Neurons were filled with Biocytin (5 mg/mL) for post-experimental identification. EPSCs were evoked using optical stimulation (473 nm; 4-6 sec.; 6-7 mW) every 60 sec, for 15 min. DNQX (25 µM) and DL-AP5 (50 µM) were applied to block AMPA and NMDA receptors. After 10 min. of drug application, EPSCs were evoked using the same procedure for another 15 min. Stimulation protocols were generated and signals acquired using Clampex 11.2 software (Molecular Devices). Series resistance was monitored and if it increased >20% during recordings, the data were discarded. Data was analyzed using WinWCP software (Courtesy of Dr. John Dempster, Strathclyde University, Glasgow, UK).

### Anatomical tracing

CRH-ires-cre mice founder animals were kindly provided by Dr. Dennis Sparta (University of Maryland, School of Medicine). Heterozygous CRF-ires-cre mice received a 200nl injection of AAV pCAG-FLEX-tdTomato-WPRE virus and wildtype micereceived a retrograde congucated cholera toxin (CTb) at a rate of 20nl/min using a 1µL Hamilton Syringe (Knurled Hub Needle, 25G) in the ventral CA1 (AP: -2; ML: ± 3.5; DV: -4.5 from the top of the skull). Four weeks after the injections, mice were perfused with 4% parafolmadehyde, and then, their brains were excised and stored in 4% parafolmadehyde for at least 24 hours prior to sectioning with a vibratome (50μm thickness). For the CTb mice, brains were fresh frozen and processed for RNAscope as described above.

### RNAscope for CRF and Fos

We utilised RNAscope Fluorescent Multiplex Detection Reagent Kit (ACDbio; catalog number: 320851) according to manufacturer’s instructions. Mice were euthanised via cervical dislocation thirty minutes following exposure to cotton-tipped swabs containing either male or female experimenters’ scent. Brains were excised and stored in -80°C. Fresh frozen brain sections (20µm) were collected using a cryostat (Leica CM1900) and directly mounted onto charged slides. Slides were fixed in 4% paraformaldehyde in 1x PBS, dehydrated in increasing concentrations of ethanol (50% 70% 100% 100%), air-dried, and outlined with a hydrophobic barrier pen. Slides then underwent Protease Plus treatment for 5 mins, followed by two 1x PBS wash for 2 mins each. Sections were hybridized with RNAscope Z probes against CRH and Fos for 2 h at 40^ᵒ^C, followed by 2 washes in 1x RNAscope Washing Buffer, 2 mins each. After hybridization, sections were incubated with AMP1 (30 min), AMP2 (15 min), AMP3 (30 min), and AMP4A (15 min) sequentially with 2 washes in 1x RNAscope Washing Buffer (2 mins each) at RT in between each steps. Lastly, sections were incubated in DAPI for 1 min at RT, rinsed, coverslipped with Vectashield Vibrance antifade mounting medium (Vector), and allowed to set overnight at 4° C prior to imaging. Analysis of RNAscope was performed by the automated image analysis software Imaris v9.2.1 (Oxford instruments). For the analysis of the CRF and Fos, a positive cell was defined as as one that expressed ≥5 CRF transcripts and ≥3 Fos trancripts. For the co-localisation analysis, a positive cell was defined as one that expressed ≥5 CRF transcripts and ≥3 Fos trancripts. Results from Imaris were validated via manual counting by an experimenter blind to the experimental groups.

### *In Vivo* Fiber photometry

Heterozygous *CRF*-ires-cre mice received 250nl bilateral infusions of retrograde AAV-syn-FLEX-jGCaMP7s-WPRE in the CA1 (AP: -2.3; ML: ±3.25; DV: -4.8 from top of the skull) in the ventral CA1. Three to four weeks after the virus infusion, mice underwent orchiectomy or sham surgeries and were bilaterally implanted with fiber optic cannulae (200μm, 0,37NA) in the EC. Following a minimum of 10-day recovery, were exposed to the male, female and control scent in the Y-maze equipment. The lenth of the arms was 11.4cm with a metal barrier to fit the tip of the swab was fitted at the end of the arm. Mice were habituated with the equipment for 8 min follwed by 8 min of test were swabs were included in each arm. During habituation and test mice were connected to the fiber photometry system (Neurophotometrics, San Diego, CA, USA) to record calcium transients in the CRF EC soma that project to the CA1 using a low-autofluorescence bifurcated optic patch cable (Doric Lenses Inc., Quebec, Canada). Independence of signals was tested prior to the start of the experiment. Excitation wavelengths used were 470 nm for the calcium-dependent rjGCaMP7s signal, temporally interleaved with 410 nm light at 40 Hz for the calcium-independent (isosbestic) signal. During the test period, photometry data was recorded timestamped using costum made code in Bonsai (San Diego, California). The fiber photometry data were analyzed offline using pMAT ^30^. Through the use of pMAT our data were corrected for photobeaching and the 410 nm signal was scaled and fitted to the 470 nm one before subtraction to correct for any motion artifacts. The Z-score was calcuated of the perievent trial, which was defined as the 5-sec prior to the entrance in each arm and 10 sec-after the entance. The signals for all the trials were averaged for each animal. Following the calculation of z-scores a smoothing filter was applied to eliminate high grequency noise and the area under curve was claculated. Each mouse was exposed to different experimenter scent and the location of the swabs was counterbalanced.

### Statistical analysis

Details of all *n* numbers. statistical analyses and results are provided in the *Supplementary Table 1*. Required samples sizes were estimated based upon on our past experience performing similar experiments. All statistical tests were two-tailed, and significance was set at *p*<0.05. Data were tested for equal variance and Gaussian distribution using the Brown-Frosythe test and Kolmogorov-Smirnov test, respectively. When normality and equal variance was achieved parametric statistical tests were performed. Correction for multiple comparisons for data analyzed with parametric ANOVAs was performed using the Holm-Sidak *post-hoc* test. When normality and/or equal variance of samples failed non-parametric tests were used. Correction for multiple comparisons for data analyzed with the non-parametric Kruskal-Wallis test was performed using the two-stage linear step-up procedure of Benjamini, Krieger and Yekutieli. Non-linear regression curve-fit was used to plot the RNAscope results using the one-phase exponential decay model followed by the extra-sum-of-squares test. Statistical analyses were performed using GraphPad Prism v8.3 except from the Wilcoxon matched-pairs signed rank tests, which were performed using SigmaPlot v14.0.

## Acknowledgements

We thank Dr. Dennis Sparta for providing us the CRF-IRES cre founder mice. We thank all the volunteers participating in these experiments.

## Author contributions

PG and TDG were responsible for the overall experimental design. TMM and KP performed the RNAscope experiments. TMM performed automated RNAscope analysis with supervision by LHT and SMC. DMG and RSD performed the independant FST replication at Yale university. CFP and CA perfused mice and processed the brains for expression and cannula implantation confirmation. DD perfomed the whole-cell electrophysiology experiment with supervision from EFP. LP and PG performed analysis for the fiber photometry experiments. CEJ and BWS performed qEEG surgeries. BWS and PG performed qEEG data analysis. CEJ performed the FST. PZ, JNH, PG, BWS, and SMC performed the PK studies (injections, euthanasia and tissue collection) and experiments utilizing single experimenters. RM and JL conducted bioanalytical quantitation of ketamine and metabolites. PY and CAZ performed the western blot experiments. SMT helped design and analyse the electrophysiological experiments. PG conducted the experiments and their analysis unless otherwise noted. PG and TDG outlined and wrote the paper, which was reviewed by all authors.

## Materials and Correspondance

The authors declare that all data supporting the findings of this study are available within the paper and its Supplementary Information files. Correspondence and requests for materials should be addressed to T.D.G.

Supplementary Table 1, Figures and Extended data figures along with Figure legends follow on the next pages

---

**Extended data Fig 1:**
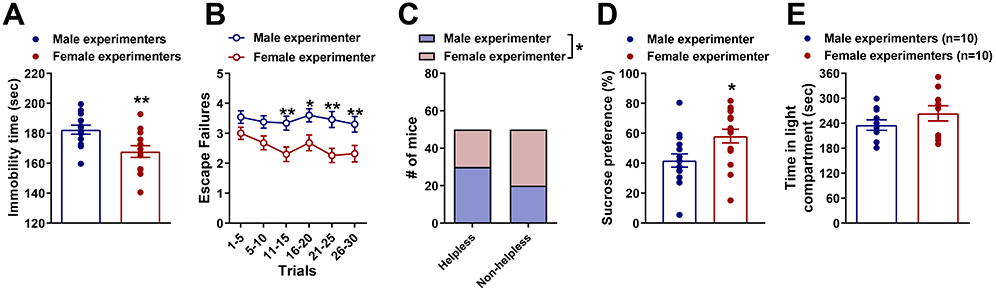
Sex of human experimenter effects on stress-related behaviours. **(A)** Immobility time measured in the forced swim test (FST) following saline injections by male and female experimenters in CD1 mice combined from all the experiments performed for the present manuscript where the mean immobility time of each experiment was used. **(B, C)** Escape failures following inescapable shock training in the learned helplessness paradigm in Swiss-Webster (CFW) mice handled by a male or a female experimenter. **(D)** Average sucrose preference over 48 hours following 10-days of chronic social defeat performed by a male and female experimenter and **(E)** time spent in light compartment in the light/dark box performed by a male and a female experimenter. Data shown are mean ± S.E.M. * *p<*0.05; ** *p<*0.01.

**Extended data Fig. 2:**
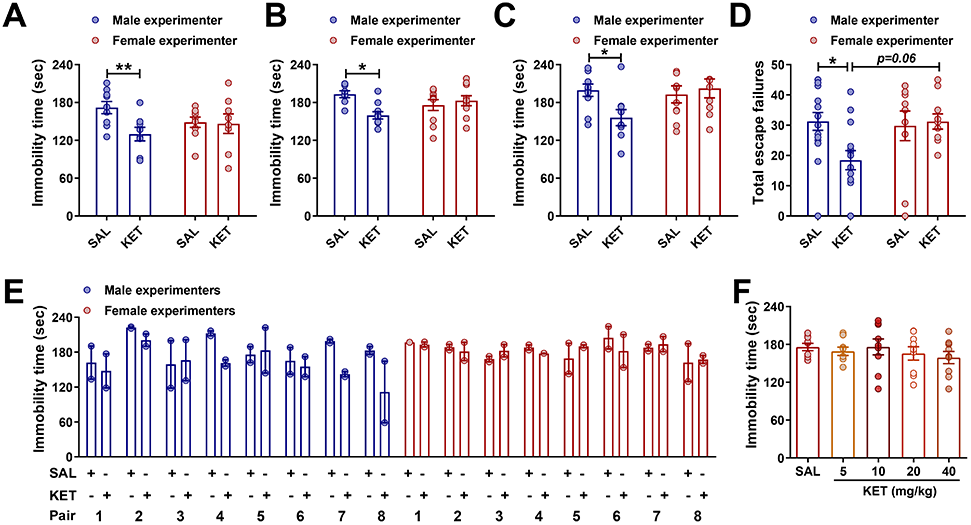
Effects of the sex of human experimenter on the antidepressant responses to ketamine. **(A)** Immibility time in the forced-swim test (FST) 24 hours post- saline (SAL; 7.5 ml/kg) and ketamine (KET; 10 mg/kg) injections by a male and female experimenter in male CD1 mice. **(B)** Immobility time in the FST 1 hour post-SAL or KET (10 mg/kg) injection by a male or female experimenter in female CD1 mice. **(C)** Immobility time 1-hour post-SAL (7.5 ml/kg ) or KET (10 mg/kg) injection by a male and female experimenter in male CD1 mice pre-exposed to a 15-min swim session 24 hours prior to the FST. **(D)** Total escape failures in the learned helplessness paradigm following SAL (7.5 ml/kg) or KET (10 mg/kg) injections by male and female experimenters in male Swiss-webster (CFW) mice. **(E)** Immobility scores in the FST 1-hour following SAL (7.5 ml/kg) or KET (10 mg/kg) injections for each individual experimenter in male CD1 mice, performed at a different institution. **(F)** KET dose-response (5, 10, 20, 40 mg/kg) in the FST 1-hr post-injection by a female experimenter in male CD1 mice. Data are shown as mean ± S.E.M. * *p<*0.05, \*\**p<*0.01

**Extended data Fig. 3:**
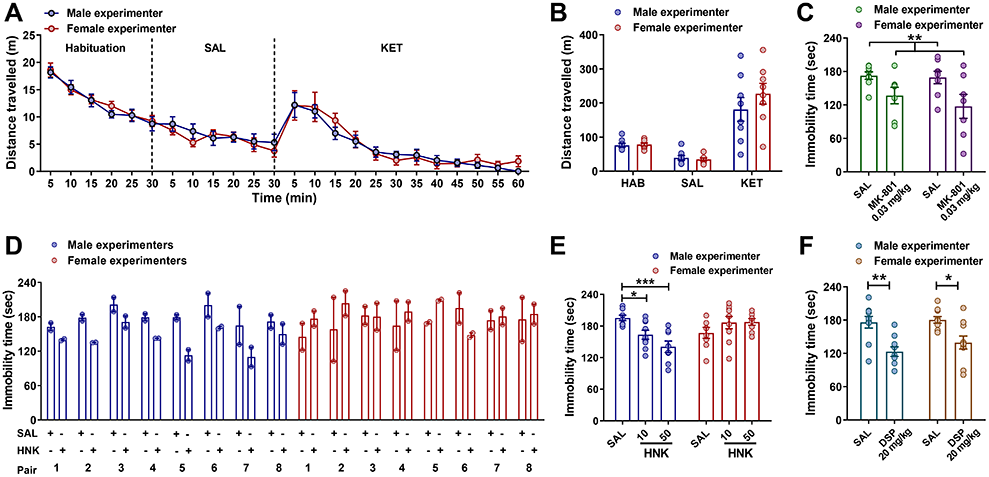
Sex of human experimenter effects on NMDAR inhibition dependent behavioural effects. **(A-B)** Distance travelled in the open-field test in male CD1 mice that received no treatment (HAB; habituation) followed by injections of saline (SAL; 7.5 ml/kg) and then ketamine (KET; 10 mg/kg) administered by a male or female experimenter. Immobility time in the forced-swim test 1-hour post-injection following administration of the **(C)** N-methyl-D-aspartate (NMDAR) receptor antagonist, MK-801 (0.03 mg/kg), **(D-E)** ketamine metabolite, (*2R,6R*)-hydroxynorketamine (HNK; 10 and 50 mg/kg), and **(F)** the classical antidepressant desipramine (DSP; 20 mg/kg) *vs* SAL (7.5 ml/kg) injections by a male and female experimenter/s. Data are shown as mean ± S.E.M. * *p<*0.05; ** *p<*0.01; *** *p<*0.001.

**Extended data Fig. 4:**
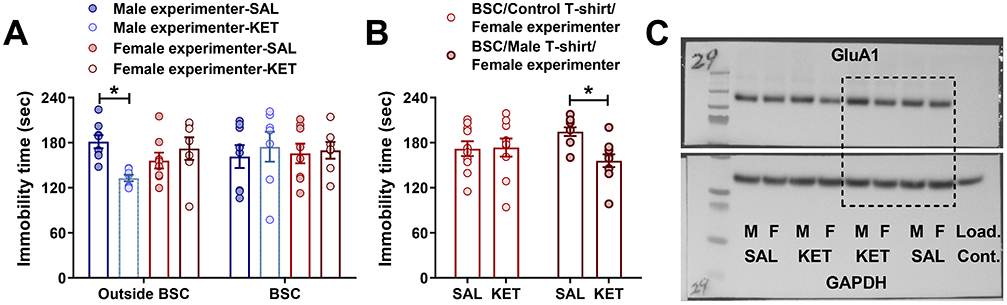
Effects of experimenter scent on the antidepressant-like responses of ketamine. **(A)** Elimination of experimenter scent by administering saline (SAL; 7.5 ml/kg) or ketamine (KET; 10 mg/kg) within a biosafety cabinet (BSC) and testing the mice in the forced swim test (FST) 1-hour post-injection. **(B)** Immobility time in mice tested in the FST 1-hour following injections of SAL (7.5 ml/kg) and KET (10 mg/kg) performed on a male worn t-shirt within the biosafety cabinet by a female experimenter (C) Whole gel images of the AMPAR receptor subunit GluA1 and GAPDH for the representative western blot images (in Fig. 2) following injection of SAL or KET by male (M) and female (F) experimenters. Data are shown as mean ± S.E.M. * *p<*0.05. Load. Cont. = loading control

**Extended data Fig 5:**
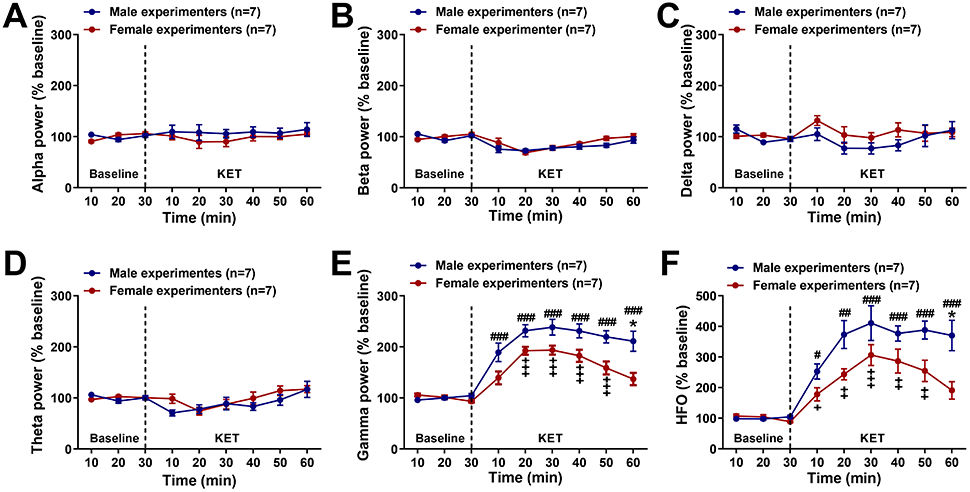
Effects of experimenter scent on the quantitative electroencephalographic oscillations. Effects of ketamine (KET; 10 mg/kg) administration by male and female experimenters on cortical quantitative electroencephalographic (qEEG) measurements using the traditionally defined frequency bands **(A)** alpha (8-12 Hz), **(B)** beta (13-29 Hz), **(C)** delta (1-4 Hz), **(D)** theta (4-8 Hz), **(E)** gamma (30-100 Hz), and **(F)** high frequency oscillations (100-160 Hz). Data are normalised to baseline, and the dashed vertical line indicates the time point of ketamine administration. Data are shown as mean ± S.E.M. *, ^+^, ^#^ *p<*0.05, ; ^++^, ^##^ *p<*0.01; ^+++, ###^ *p<*0.001. Differences between ketamine response administered by male and female experimenters is indicated by *. Differences between baseline and ketamine is indicated by + for male experimenters and by # for female experimenters.

**Extended data Fig. 6:**
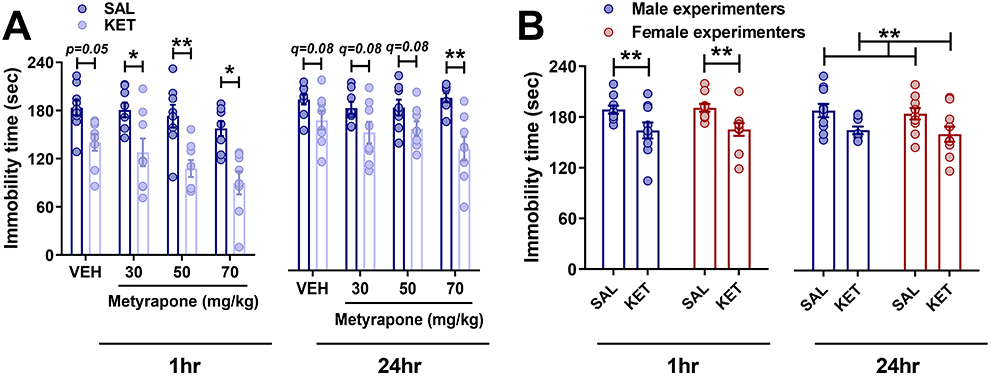
Effects of corticosterone on the antidepressant responses to ketamine. **(A)** Forced-swim test (FST) immobility measured in mice that received metyrapone (30, 50 and 70 mg/kg) prior to KET (10 mg/kg) and tested 1- and 24-hours later. **(B)** Immobility time measured in the FST following saline (SAL; 7.5 ml/kg) or ketamine (KET; 10 mg/kg) administration by male and female experimenters to male BALB/c mice. Data are shown as mean ± S.E.M. * *p<*0.05; ** *p<*0.01; for non-parametric analysis ** *q<*0.01

**Extended data Fig. 7:**
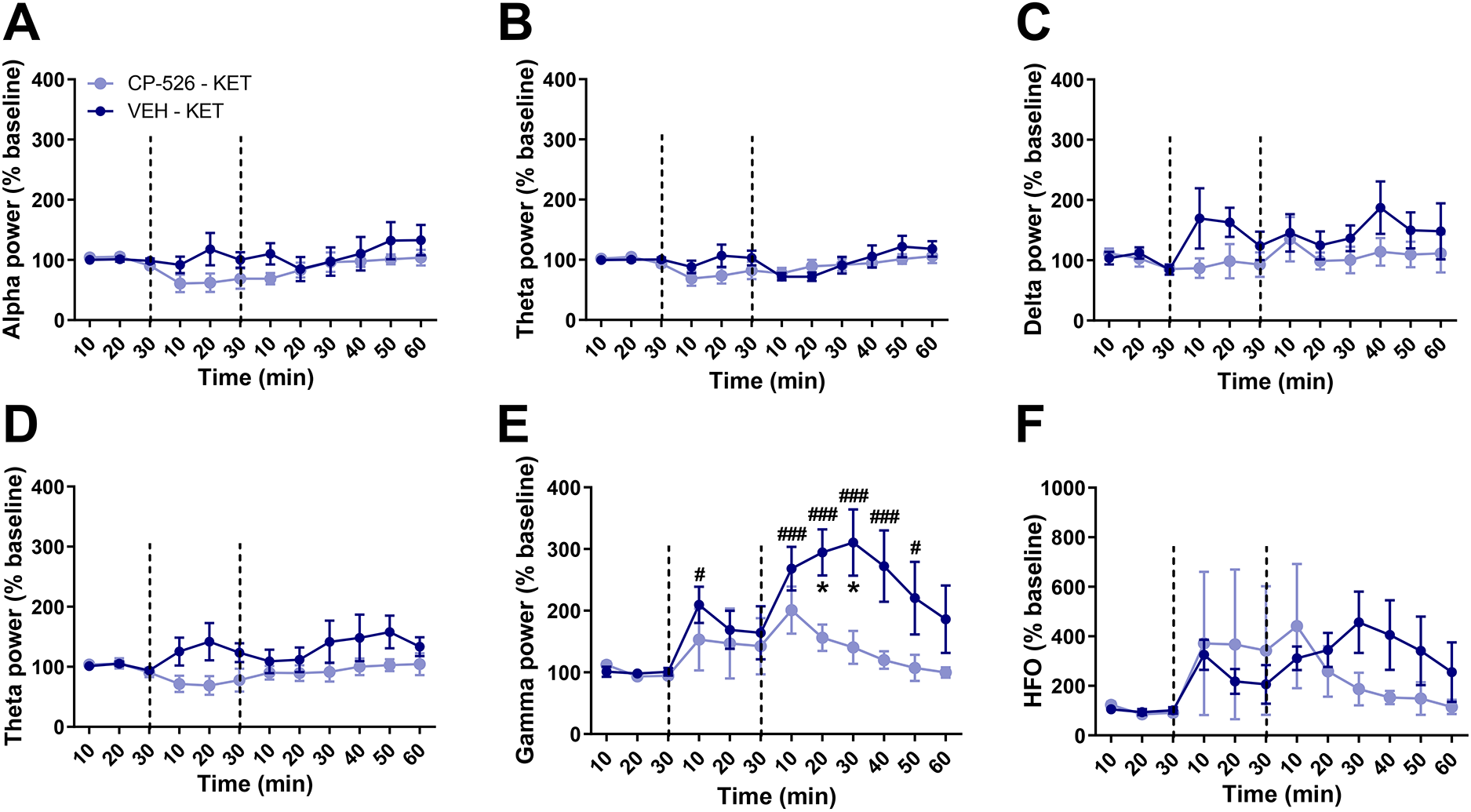
Effects of the CRF1 antagonist, CP-154,526, on electroencephalographic measures following ketamine administration. Effects of the corticotropin-releasing factor 1 antagonist (CRF1), CP-154,526 (CP-526; 30 mg/kg) or vehicle (VEH; 1.5 ml/kg) prior to ketamine (KET; 10 mg/kg) administration by a male experimenter on cortical qEEG mesearuments using the traditionally defined frequency bands **(A)** alpha (8-12 Hz), **(B)** beta (13-29 Hz), **(C)** delta (1-3Hz), **(D)** theta powers (4-8 Hz), **(E)** gamma (30-100 Hz), and **(F)** high frequency oscillations (100-160 Hz). Data are normalised to baseline. The first dashed vertical line indicates the time point of VEH or CP-526 administration and the sec dashed vertical line indicates the time point of KET administration. Data are shown as mean ± S.E.M. *, ^#^ *p<*0.05, ; ^###^ *p<*0.001. Differences between mice pre-treated with CP-526 and VEH are indicated by *. Differences between the baseline and treatment in mice that received VEH prior to KET are indicated with #.

**Extended data Fig. 8:**
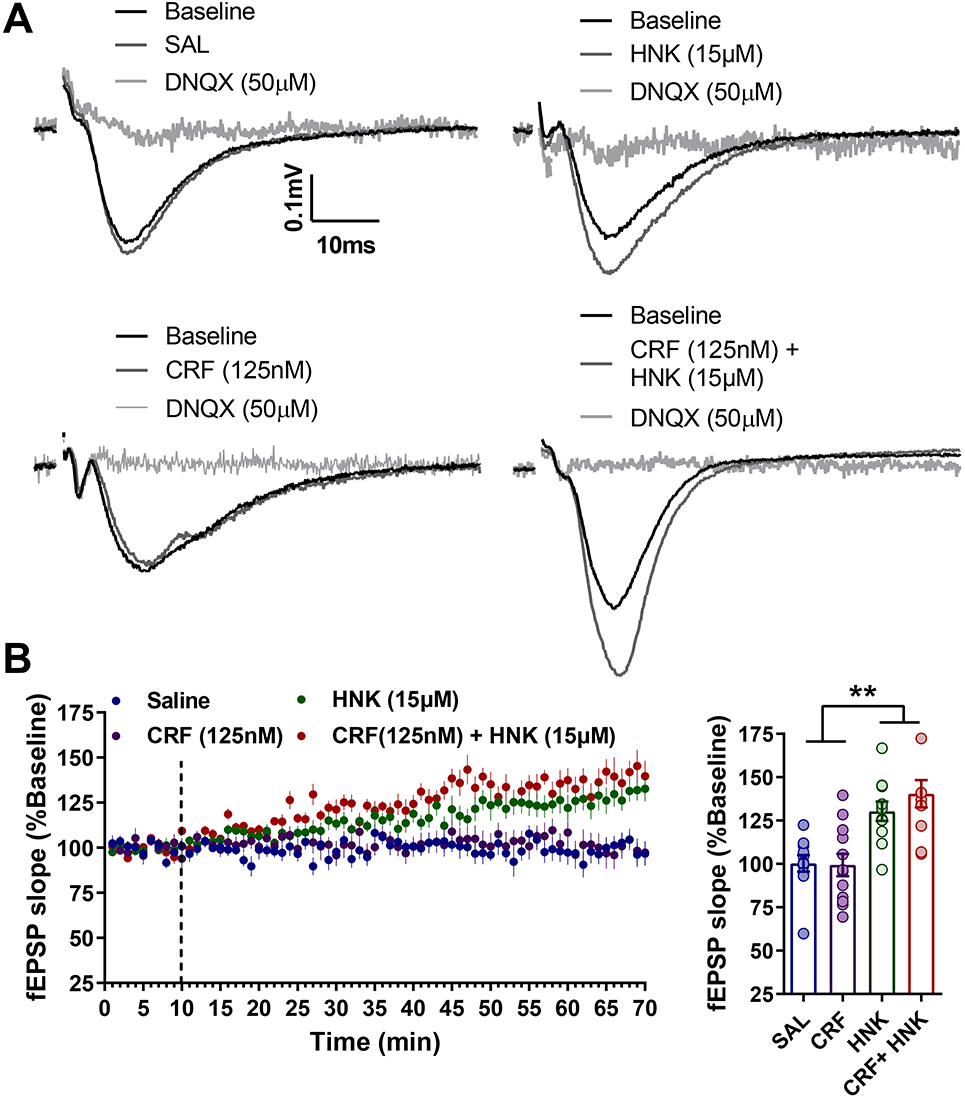
Effects of the combined CRF and (*2R,6R*)-hydroxynorketamine on field excitatory postsynaptic potentials in hippocampal slices in the SC-CA1 pathway. **(A)** Representative traces of Schaffer collateral-hippocambal CA1 (SC-CA1) field excitatory postsynaptic potentials (fEPSPs) and **(B)** quantification of fEPSPs slopes following SC-CA1 pathway stimulation during wash-in with Saline (SAL), coricotropin-rekeasing factor (CRF; 125nM), (*2R,6R*)-hydroxynorketamine (HNK) (15μM) or combined CRF (125nM) + (*2R,6R*)-HNK (15μM). Data are shown as mean ± S.E.M. ** *p<*0.01.

**Extended data Fig. 9:**
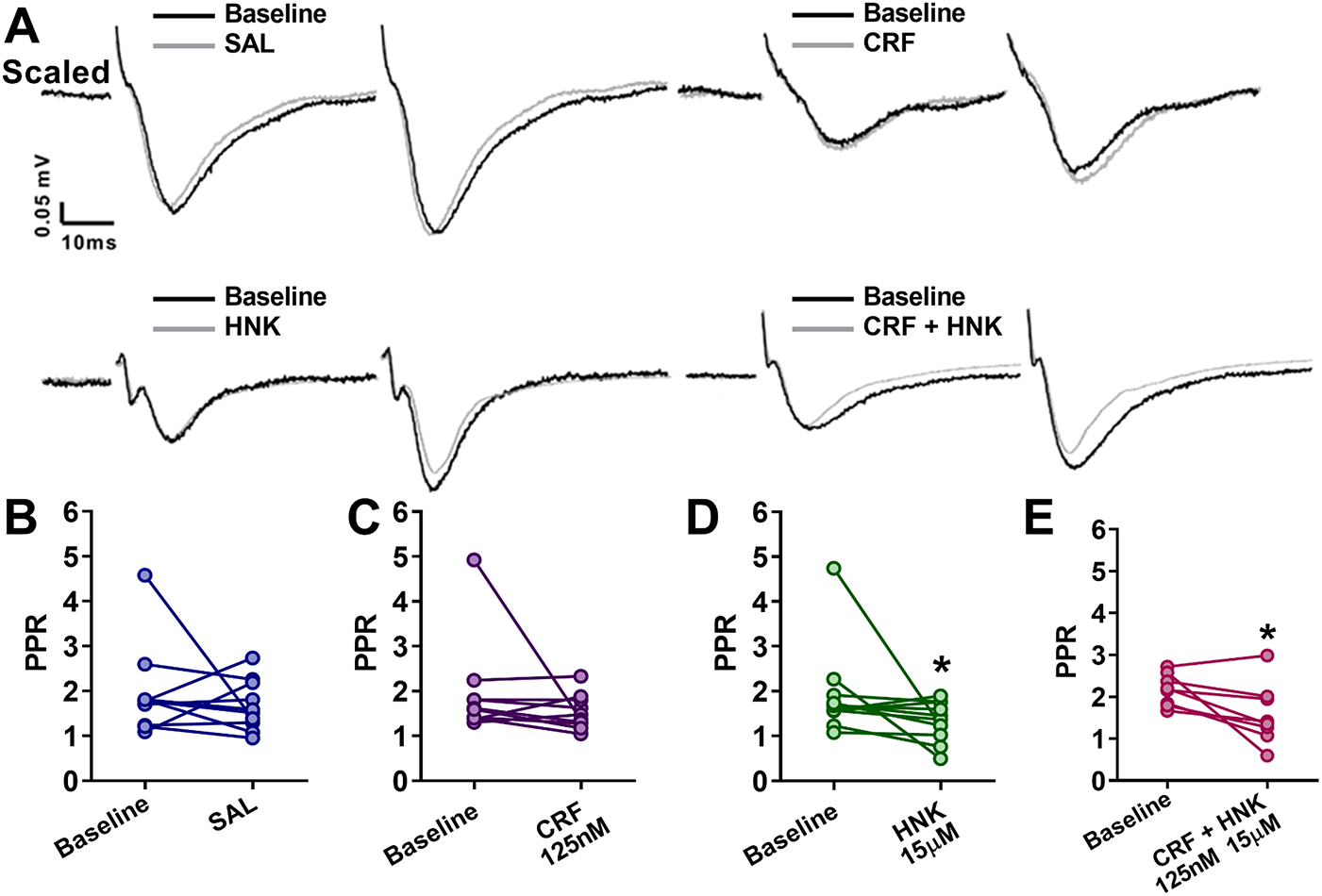
Effects of the combined CRF and (*2R,6R*)-hydroxynorketamine on paired-pulse ratio in hippocampal slices in the SC-CA1 pathway. **(A)** Representative traces and quantification of paired pulse-ratio (PPR) following stimulation of the Schaffer collateral-hippocampal CA1 (SC-CA1) pathway pre- and post wash-in with **(B)** saline (SAL), **(C)** corticotropin-releasing factor (CRF; 125 nM), **(D)** (*2R,6R*)-hydroxynorketamine (HNK) (15μM) and **(E)** CRF (125nM) + (*2R,6R*)-HNK (15μM). * *p<*0.05.

**Extended data Fig. 10:**
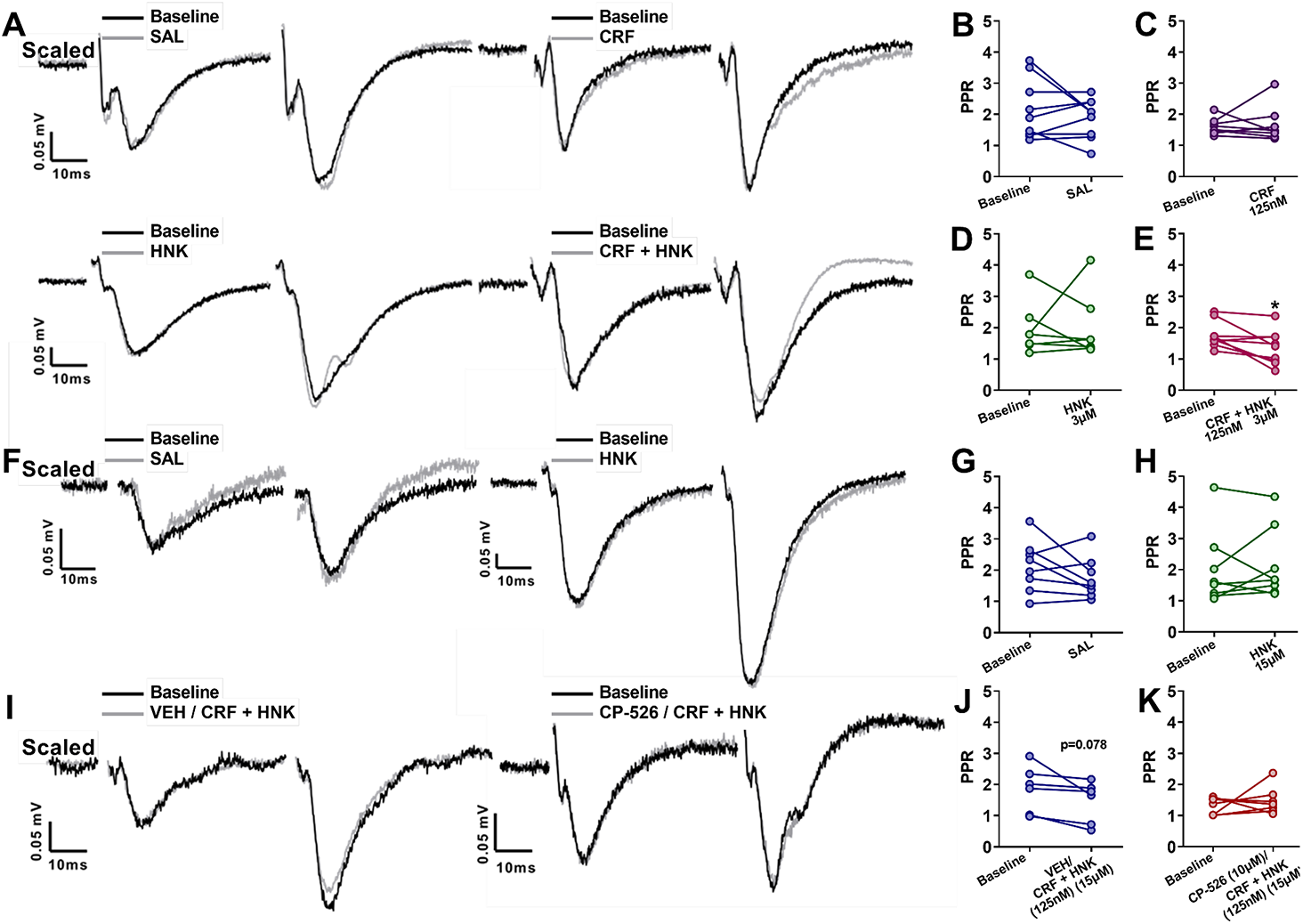
Effects of the combined CRF and (*2R,6R*)-hydroxynorketamine on paired-pulse ratio in the EC-CA1 pathway. **(A)** Representative traces and quantification of paired-pulse ratio (PPR) following stimulation of the entorhinal cortex-hippocampal CA1 (EC-CA1) pathway pre- and post wash-in with **(B) s**aline (SAL), **(C)** corticotropin-releasing factor (CRF;125 nM), **(D)** (*2R,6R*)-hydroxynorketamine (HNK) (3μM) and **(E)** CRF (125nM) + (*2R,6R*)-HNK (3μM). **(F)** Representative traces and quantification of paired-pulse ratio following stimulation of the EC-CA1 pathway pre- and post wash-in with **(G)** SAL and **(H)** (*2R,6R*)-HNK (15μM). **(I)** Representative traces and quantification of PPR from the EC-CA1 stimulation pathway pre- and post wash-in with **(J)** Vehicle (VEH) and CRF (125nM) + (*2R,6R*)-HNK (3μM) and **(K)** CP,154-526 and CRF (125nM) + (*2R,6R*)-HNK (3μM). * *p<*0.05.

**Extended data Fig 11:**
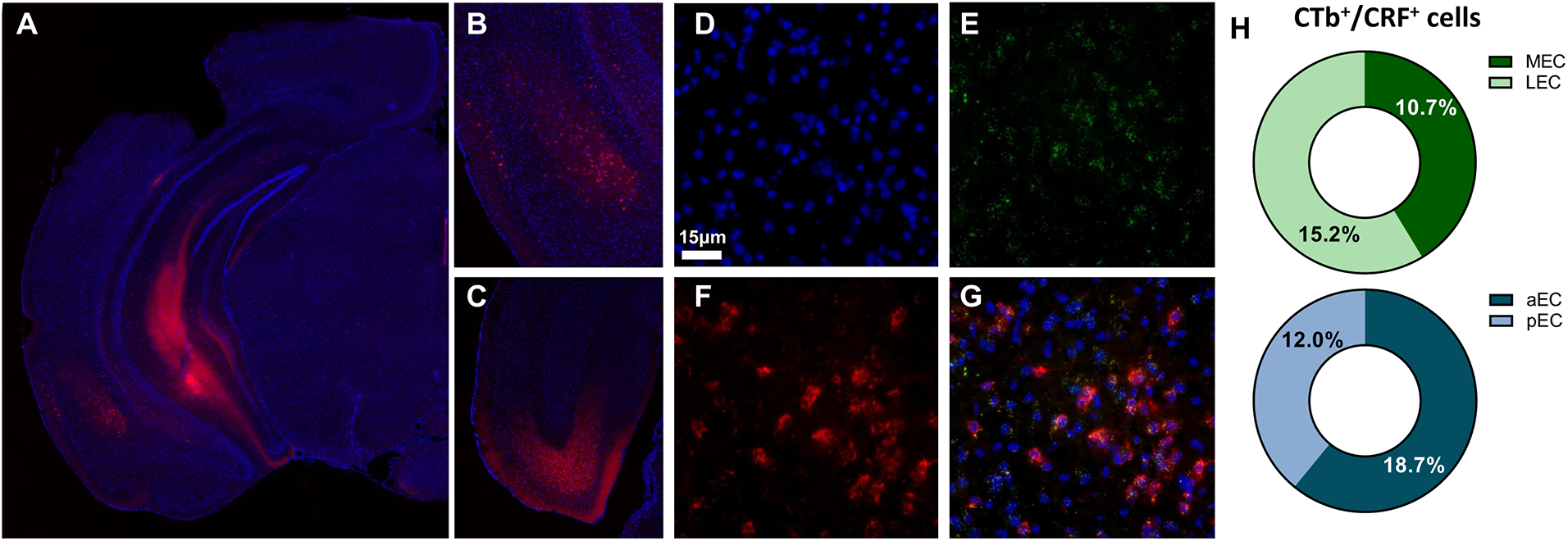
Identification of EC to CA1 projectionS. **(A)** Representative images of the injection site at ventral CA1 with retrograde congugated cholera toxin and the **(B,C)** labelling in the anterior and posterion entorhinal cortex (EC). Representative RNAscope images from the EC revealing **(D)** DAPI labeling, **(E)** CRF trancripts labelling, **(F)** CTb labelling and **(G)** the collabelling between the tracer and CRF trancripts. **(H)** Quantification of RNA scope and tracer colabelling at the anterior (aEC) and posterior EC (pEC) and the lateral (LEC) and medial EC (MEC)

